# *DifferentialRegulation*: a Bayesian hierarchical approach to identify differentially regulated genes

**DOI:** 10.1101/2023.08.17.553679

**Authors:** Simone Tiberi, Joël Meili, Peiying Cai, Charlotte Soneson, Dongze He, Hirak Sarkar, Alejandra Avalos-Pacheco, Rob Patro, Mark D Robinson

## Abstract

**Motivation:** Although transcriptomics data is typically used to analyse mature spliced mRNA, recent attention has focused on jointly investigating spliced and unspliced (or precursor-) mRNA, which can be used to study gene regulation and changes in gene expression production. Nonetheless, most methods for spliced/unspliced inference (such as RNA velocity tools) focus on individual samples, and rarely allow comparisons between groups of samples (e.g., healthy *vs*. diseased). Furthermore, this kind of inference is challenging, because spliced and unspliced mRNA abundance is characterized by a high degree of quantification uncertainty, due to the prevalence of multi-mapping reads, i.e., reads compatible with multiple transcripts (or genes), and/or with both their spliced and unspliced versions.

**Results:** Here, we present *DifferentialRegulation*, a Bayesian hierarchical method to discover changes between experimental conditions with respect to the relative abundance of unspliced mRNA (over the total mRNA). We model the quantification uncertainty via a latent variable approach, where reads are allocated to their gene/transcript of origin, and to the respective splice version. We designed several benchmarks where our approach shows good performance, in terms of sensitivity and error control, versus state-of-the-art competitors. Importantly, our tool is flexible, and works with both bulk and single-cell RNA-sequencing data.

**Availability and implementation:** *DifferentialRegulation* is distributed as a Bioconductor R package.

## 1 Introduction

Bulk and single-cell RNA-sequencing (RNA-seq) data enable estimating the abundance of both (mature) spliced (*s*) and unspliced (*u*) (or precursor) mRNA. These splicing dynamics have been previously studied from bulk data (Zeisel et al., 2011; Gaidatzis et al., 2015); furthermore, in single-cell RNA-seq (scRNA-seq) data, they have been further exploited by RNA velocity tools, that infer the time derivative of the gene expression state of cells (La Manno et al., 2018; Bergen et al., 2020). In these approaches, *s* and *u* abundances are compared to their (estimated) equilibrium values. Intuitively, if a gene has a higher relative abundance of *u* reads than at steady-state, in the near future the *s* mRNA will increase because we expect a higher abundance of newly spliced mRNA, compared to the amount of *s* mRNA that is going to be degraded (Figure 1). Therefore, gene expression (i.e., *s*) is currently increasing, and we can think of this gene as being up-regulated. Conversely, if the relative abundance of *u* reads is lower than at its equilibrium, in the near future, the amount of *s* mRNA will decrease, because the newly spliced mRNA will not fully compensate for the degraded mRNA (Figure 1). In this case, gene expression is currently decreasing; hence, we can conclude that the gene is being down-regulated.

**Figure 1:**
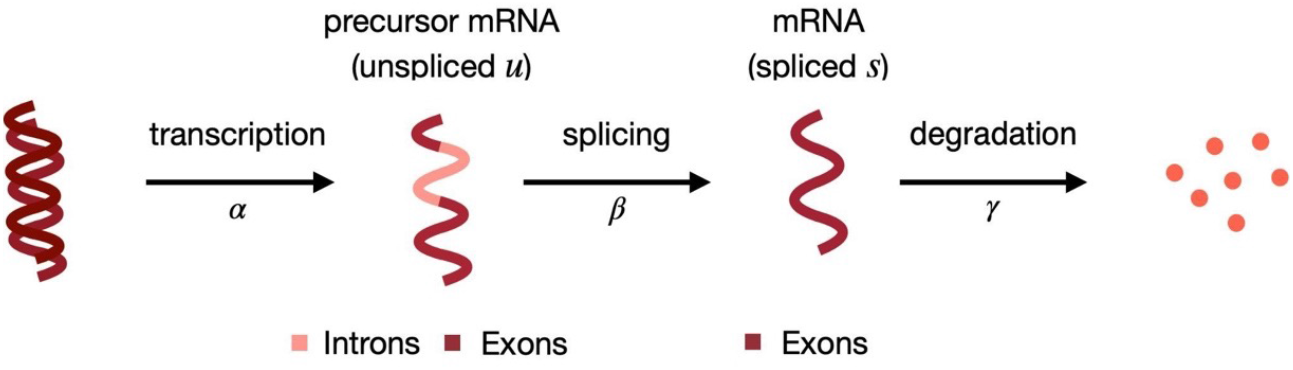
Splicing dynamics (Weiler et al., 2022): unspliced mRNA (*u*), containing both introns and exons, is transcribed from DNA (at rate *α*); then, splicing (at rate *β*) leads to spliced mRNA (*s*), which is eventually degraded (at rate *γ*).

Here, following a similar rationale, we aim at identifying differences in gene regulation between experimental conditions (e.g., treatments), by comparing the relative abundance of *u* reads, denoted by *π*_*U*_. In particular, for a given gene, if *π*_*U*_ is higher in condition *A* than *B*, we speculate that the gene is being up-regulated in *A*, compared to *B*. Note that this is different from canonical differential gene expression tests, which focus on differences in the overall abundance of *s* reads. Instead, our goal is to identify differences in the direction that gene expression is currently undergoing. In particular, it was found that unspliced mRNA peaks, on average, 15 minutes before spliced mRNA, and can be taken as a proxy for nascent transcription (Hendriks et al., 2014). Furthermore, changes in unspliced mRNA are thought to be indicative of changes in post-transcriptional regulation (Gaidatzis et al., 2015). Therefore, identifying variations in *π*_*U*_ can provide valuable insight into gene regulation changes between conditions.

From a technical point of view, RNA-seq data is characterized by a large degree of quantification uncertainty due to multi-mapping reads, i.e., reads compatible with multiple transcripts or genes (McDermaid et al., 2018; Dharshini et al., 2020). Furthermore, when analyzing splicing dynamics, we consider both splice versions of each gene/transcript; this doubles the number of transcripts (bulk data) or genes (single-cell data) in the reference, and increases even more mapping ambiguity.

Two approaches, namely *eisaR* (Gaidatzis et al., 2015) and *BRIE2* (Huang and Sanguinetti, 2021), have been proposed to compare splicing dynamics between groups of samples from bulk and scRNA-seq data, respectively. The first approach uses the *edgeR* (Robinson et al., 2009) differential pipeline, based on a negative binomial distribution, where samples and groups are used as covariates for the mean parameter. The second method, instead, implements a Bayesian regression approach on percent spliced-in values, with samples and groups modelled as covariates; however, this approach was found to be extremely computationally intensive.

Additionally, other tools, originally designed to detect differences in alternative splicing patterns, could also be employed to discover changes among *s* and *u* reads. Notably, *DRIMSeq* (Nowicka and Robinson, 2016), *satuRn* (which has two variants: one for bulk and one for single-cell data) (Gilis et al., 2022), *SUPPA2* (Trincado et al., 2018), and *DEXSeq* (Anders et al., 2012) performed well in recent benchmarks (Love et al., 2018; Tiberi and Robinson, 2020; Gilis et al., 2022). In particular, *DEXSeq*, for the purpose of our analyses, could be applied to transcript estimated abundance (Love et al., 2018) (referred to throughout as *DEXSeq_TECs*), or to equivalence classes counts (Cmero et al., 2019) (denoted by *DEXSeq_ECs*), where equivalence classes (ECs) are collections of reads compatible with the same set of transcripts (including splicing status). Such ECs, and their multiplicities, are typically used to model the variability of multi-mapping reads. The majority of differential methods, in our case, *eisaR, DRIMSeq, satuRn, SUPPA2* and *DEXSeq_TECs*, input estimated counts, and thus fail to account for the noise in those estimates. Conversely, *DEXSeq_ECs* avoids this issue by performing differential testing directly on equivalence classes. However, while this approach accounts for reads mapping between *s* and *u* versions of a transcript, it does not handle reads mapping to multiple transcripts, which are discarded, hence resulting in a loss of data. Moreover, while most methods presented above can test genes or transcripts directly, *SUPPA2* and *DEXSeq_ECs* perform differential testing on exon junctions and ECs, respectively, which results in multiple statistical tests for each transcript, that are then aggregated to the transcript level. When applied to scRNA-seq data, *BRIE2, DRIMSeq, satuRn*, and *DEXSeq_TECs* can partially account for the quantification uncertainty, by treating separately ambiguously mapping reads (i.e., those mapping to both *s* and *u* versions of a transcript); such reads are denoted by *a*. However, the ambiguity in multi-gene mapping reads cannot be modelled.

To overcome these challenges, we propose a Bayesian approach that accounts for the quantification uncertainty via a latent variable model, and allocates reads to their transcript or gene of origin, and corresponding splice version. Our approach also allows for sharing of information across samples, via a hierarchical framework, and genes, via informative priors. We designed a double analysis framework, based on two distinct *ad hoc* algorithms to analyze bulk and single-cell RNA-seq data, accounting for the specifics of each type of data. In particular, bulk protocols enable studying transcript-level signals across all cells, while single-cell data offer high cellular resolution but do not allow accurate transcript-level inference. Here, we take advantage of the information that each offers: our bulk approach targets changes at the transcript level (across all cells), while our single-cell method identifies cell-type specific changes (at the gene level), e.g., genes that are differential in a cell type but not in others. Below, we illustrate both approaches (Section 2), describe our benchmarks (Section 3), and discuss results (Section 4).

## 2 Methods

### 2.1 Model for bulk data

*DifferentialRegulation* takes as input the equivalence classes counts derived from RNA-seq reads, and recovers the overall abundance of each transcript. Assume that, for a given experimental condition, we collect RNA-seq data for *N* samples (i.e., biological replicates), with a total of *T* transcripts; we define by 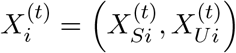 the vector indicating the overall abundance of spliced and unspliced reads coming from transcript *t* in sample *i*, with *t* = 1, …, *T*, and *i* = 1, …, *N*. Our approach is built around two models.

The first one is a multinomial model for the abundance of reads across the *T* transcripts:

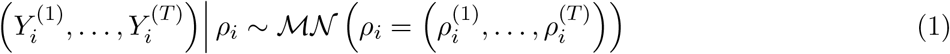

where 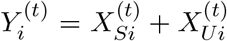 is the overall abundance (aggregated across both splice versions) of transcript *t* in sample *i*, and 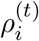 indicates the relative abundance for the *t*-th transcript in the *i*-th sample, with 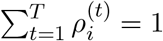.

The second model is a hierarchical beta-binomial distribution for the abundance of reads within the *s* and *u* versions of each transcript:

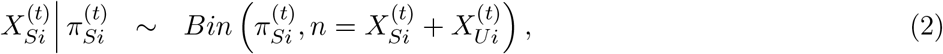

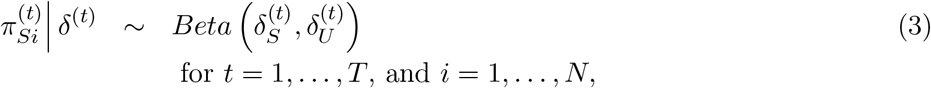

where 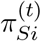 represents the relative abundance of spliced reads for transcript *t* in the *i*-th sample, 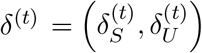 represent the hyper-parameters of the hierarchical model, and *Beta*(*a, b*) indicates the beta distribution with mean 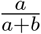 and variance 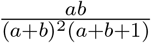. Note that, for easier interpretation, the hyperparameters can be reparametrized as 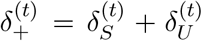, usually referred to as the precision parameter, which indicates the sample-to-sample variability, and 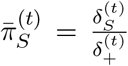(or 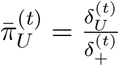), denoting the group-level relative abundance of *s* (or *u*) reads for transcript *t*, respectively. Note that we chose not to use a hierarchical prior for *ρ* for two main reasons: i) overall transcript abundances are easier to infer (hence the benefit of sharing information across samples is smaller), and ii) we wanted to limit the model complexity, and its computational cost.

Since the values of *X* (and hence *Y*) are not observed, they are treated as latent states and are sampled from their conditional distribution (see Section 2.3). In particular, we allocate multi-mapping reads among the transcript(s) and respective splice version(s) they are compatible with. For instance, in sample *i*, consider a read compatible with the *s* version of transcript *w*, and the *u* version of transcript *z*; this read will be allocated to the former and latter cases with probability proportional to 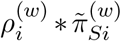, and 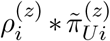, respectively, where 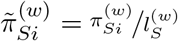 and 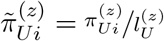, with 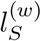 and 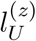 being the effective lengths of the *s* version of transcript *w* and the *u* version of transcript *z*, respectively. Normalizing for the transcript effective lengths ensures that the probability of allocating multi-mapping reads does not depend on how long transcripts are, and was previously found to improve model accuracy (Soneson et al., 2015; Tiberi and Robinson, 2020).

### 2.2 Model for single-cell data

Our framework for single-cell data is similar to the bulk approach, but presents four key differences, as summarized below. First, droplet scRNA-seq data have little resolution at the transcript level; therefore, analyses are performed on genes instead of transcripts. Second, cells are typically clustered (usually in cell types): when cell clusters are available, we separately analyze each cluster, and identify cluster-specific changes in regulation. Third, data refer to individual cells: after clustering them, we use a pseudo-bulk approach and, for each sample, compute the total *s* and *u* counts across all cells in a given cluster. Fourth, reads ambiguously mapping to both *s* and *u* versions of a gene (i.e., *a* reads) cannot easily be allocated to their splice version of origin. This is because the allocation step requires the estimated probability that an ambiguous read is spliced, which cannot be accurately computed, because it depends on unknown factors (Supplementary Details). Therefore, we only use a latent variable approach for reads mapping to multiple genes; instead, *a* reads are treated separately from *s* and *u*.

Consider one experimental condition and a single cell cluster, with scRNA-seq data available for *N* samples and *G* genes; we denote by 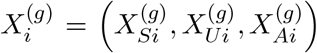 the vector with the overall abundance of spliced, unspliced and ambiguous reads (across all cells in the cluster) coming from gene *g* in sample *i*, with *g* = 1, …, *G*, and *i* = 1, …, *N*. Again, we use two models; the first one is a multinomial distribution for the abundance of reads across the *G* genes:

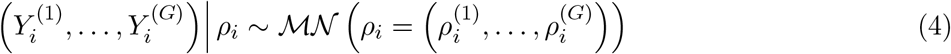

where 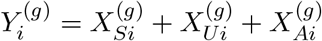 is the overall abundance (across all splice versions) of gene *g* in sample *i*, and 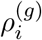 is the relative abundance for gene *g* in the *i*-th sample, with 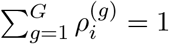.

The second model is a hierarchical Dirichlet-multinomial, which is a generalization of the beta-binomial model in (3)-(6), for the abundance of *s, u* and *a* reads within each gene:

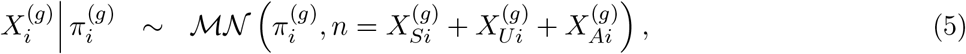

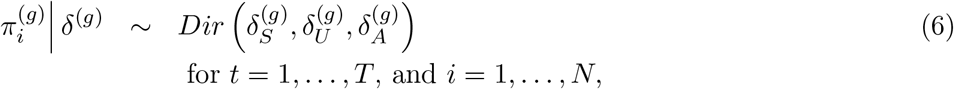

where 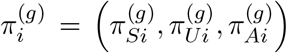 is the vector with the relative abundance of spliced, unspliced and ambiguous reads for gene *g* in the *i*-th sample, and 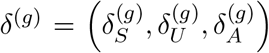 represent the hyper-parameters of the hierarchical model; Again, from the hyper-parameters, we can obtain the precision parameter 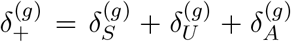, governing the sample-to-sample variability, and the group-level relative abundances of *s, u* and *a* reads for transcript 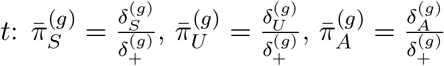, respectively.

As before, the counts in *X* are not observed and treated as latent variables, which are sampled based on *ρ*’s and *π*’s, by allocating reads to their gene of origin and respective splice version (i.e., *s, u* or *a*) (see Section 2.3). As an example, assume that, in sample *i*, a read is compatible with the *s* version of gene *w*, the *u* version of gene *z*, and the *a* version of gene *q*; this read will be allocated to the one of three cases with probability proportional to 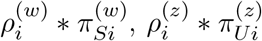, and 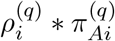, respectively. Note that, unlike in the bulk model, here we do not normalize for the effective lengths of genes; this is primarily due to two reasons. First, the effective length of genes is defined as a weighted average of the effective lengths of transcripts, weighted by transcript relative abundance, which is not known. Second, normalizing for the effective length of transcripts in the bulk model is based on the assumption that, given the same mRNA abundance, longer transcripts will produce more RNA-seq reads; however, this assumption is not always valid in scRNA-seq protocols.

### 2.3 Parameter inference

In both models, our hierarchical framework allows sharing of information between samples; we further share information across transcripts (bulk model) and genes (single-cell model) via an empirical Bayes approach. In particular, our hyper-parameters *δ*, in (3) and (6), are initially estimated from a random selection of 1,000 genes/transcripts, via *DRIMSeq* (Nowicka and Robinson, 2016); these estimates are used to formulate informative priors for all hyper-parameters (Supplementary Details). Note that, our empirical Bayes approach is very mild because each gene/transcript contributes in a tiny fraction to the prior formulation. Additionally, we assume a weakly informative conjugate Dirichlet prior distribution for *ρ*, which results in a conjugate Dirichlet posterior distribution (Supplementary Details).

If the values of *X* (and hence of *Y*) were observed, it would be straightforward to formulate the likelihood of the model in terms of the multinomial and binomial densities in (1)-(2) (bulk model), and in (4)-(5) (single-cell model). However, since *X* is not observed, in both models the likelihood of the data is defined with respect to the actual observations, which, in our case, are the number of counts in each equivalence class. In general terms, we define *θ, X* and *Z* as the objects containing all model parameters, all latent variables, and the data from all equivalence classes, respectively. The likelihood of the model, *L*(*θ*|*Z*), can be expressed as the integral over the latent states: *L*(*θ*|*Z*) = *∫*_*X*_ *p*(*Z, X* = *x*|*θ*)*dx*. Here, we use a Bayesian data augmentation approach (Tanner and Wong, 1987; Gelfand and Smith, 1990) which, instead of working with this integral, alternately samples parameters and latent states from their conditional distributions: *p*(*θ*|*Z, X*), and *p*(*X*|*Z, θ*). In particular, our sampling scheme follows a Metropolis-within-Gibbs Markov chain Monte Carlo (MCMC) algorithm where parameters are updated, in four steps, from the following conditional distributions: *δ*|*π, π*|*X, δ, ρ*|*X*, and *X*|*Z, π*. Importantly, although our approach involves many parameters, the vast majority of them are updated using a Gibbs sampler, which results in better mixing and convergence; only the hyper-parameters *δ* are sampled according to a Metropolis step, where values are proposed based on an adaptive random walk (Haario et al., 2001). Supplementary Details report, for each parameter, the prior and conditional distributions, and the sampling scheme used. Since the sampling of the latent variables is the most computationally intensive step of our algorithms, we employ an undersampling scheme where latent variables are updated every 10 iterations (users can decrease this parameter). In our benchmarks, this led to a reduction of the runtime of our full pipeline of 74% and 35%, for the bulk and single-cell models, respectively. By default, the MCMC is run for 2,000 iterations, with a burn-in of 500 iterations (parameters can be increased by users). To ensure convergence, a Heidelberger and Welch stationarity test (Heidelberger and Welch, 1983) is performed on the marginal log-posterior density of the hyper-parameters, i.e., log(*p*(*δ*|*π*)). If the test fails, the burn-in is automatically increased up to half of the chain length; if convergence is still not reached, a new chain is run, with doubled burn-in and number of iterations. Additionally, our software allows users to visually investigate convergence and mixing, by plotting (via the *plot_traceplot* function), in each group, the posterior distribution of 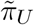, which is the key parameter of the model (see Section 2.4).

### 2.4 Comparing groups

Until now, we have shown how we infer model parameters, separately, in each group of samples; in what follows, we will describe how results across conditions are compared. To this aim, we introduce a new parameter, 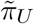 in the bulk model, we set 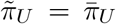, while in the single-cell approach we also account for 50% of ambiguous reads, and define 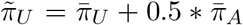. Note that, for simplicity, we have dropped gene and transcript indices from the notation. Given two groups of samples, *A* and *B*, to identify differentially regulated genes/transcripts, we compare 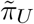 between *A* and *B*, that we call 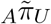 and 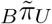, respectively. We therefore define the probability that group *B* is up-regulated, compared to group *A*, as 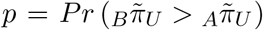, which can be easily estimated from the posterior chains. In the Results Section, we used this probability to rank genes/transcripts for *DifferentialRegulation*; in particular, we rank them according to *max*(*p*, 1 − *p*). In other words, results with *p* close to 0 or 1 are ranked first (i.e., 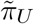 differs between groups), while results with *p* ≃ 0.5 are ranked last (i.e., 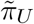 similar across groups).

In the single-cell model, we acknowledge that setting 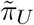 as 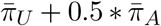 is based on the arbitrary choice of equally assigning 50% of ambiguous reads to *s* and *u*. Therefore, we also provide an alternative way to rank genes, which does not require assigning ambiguous reads. In particular, we approximate the posterior distribution of the original (*π*_*S*_, *π*_*U*_) with a bivariate normal around its posterior mode, and perform a bivariate Wald test (Li et al., 1991) to verify if *s* and *u* proportions are equivalent across groups (Supplementary Details); note that *π*_*A*_ is not considered, because it is uniquely defined by *π*_*S*_ and *π*_*U*_. Genes are then ranked based on the p-value of this test. Below, results based on the probability *p* are referred to as *DifferentialRegulation*, while those based on the Wald test are called *DifferentialRegulation_Wald*.

### 2.5 Simulation design

We designed several simulation studies to benchmark our method and competing approaches. To generate realistic simulations, we started with real datasets as anchor data. In the bulk simulation, which is an extension of the human simulation framework used in Soneson et al. (2016), we used a sample from Trapnell et al. (2013) (SRR493366) to infer the relative abundance of each transcript and splice version. Estimates of sample-to-sample variability were obtained using data from Cheung et al. (2010) and Pickrell et al. (2010), as previously described (Soneson and Delorenzi, 2013). We then used these parameters to simulate counts for each transcript (and splice version) for 6 samples, which were randomly separated in two groups. We randomly selected 2,000 transcripts as differentially regulated (DR); for each one, we inverted their *s* and *u* abundances in one of the two groups. In order to introduce quantification uncertainty in our simulation, we provided the vectors with desired transcript per million values for each transcript to *RSEM* (Li and Dewey, 2011) to simulate reads, and then mapped these reads to a reference transcriptome with *salmon* (Patro et al., 2017).

In the single-cell simulation, we started from the mouse data from Park et al. (2018), consisting of four biological replicates; in this case, however, we did not simulate counts: instead, we used estimated spliced and unspliced counts directly. We annotated cell types via *SingleR* (Aran et al., 2019), and kept the three most abundant ones. As above, we separated samples in two groups, and introduced a DR effect, separately for each cell-type, in 20% of genes, by inverting *s* and *u* counts in one of the two groups (selected at random). In order to generate cell-type-specific changes, we randomly selected distinct differential genes in each cell type. To introduce quantification uncertainty, we provided the count matrices (generated above) to a read-level simulator, *minnow* (Sarkar et al., 2019), and aligned the simulated reads via *alevin-fry* (He et al., 2022).

In both bulk and single-cell analyses, we designed several additional simulations to investigate robustness of resutls across various scenarios. First, we generated three simulations, where we added differential gene expression (DGE) between groups, with an average fold change of 3, 6 or 9. Second, in the bulk analyses only, we simulated differential alternative splicing (DAS) across conditions; DAS was not simulated in single-cell data, because it requires transcript-level resolution, which is not available in scRNA-seq protocols. Here, our aim is to identify DR genes, while DGE and DAS are nuisance effects, that we do not wish to detect. Below, we refer to DR, DR + DGE (average fold change of 3), and DR + DAS as our main simulations. Third, in the single-cell analyses only, we simulated three datasets (with DR only) varying the number of cells with zero expression (90, 95 and 99%), while keeping the overall gene abundance unchanged. Fourth, we investigated the impact of batch effects on inference. For what concerns our analyses, batch effects could potentially introduce differences in overall gene expression (DGE), in alternative splicing (DAS), or in the relative abundance of spliced reads within genes/transcripts (DR). While the impact of DGE and DAS on results can be assessed from the simulations described above, we designed a further simulation where DR is introduced across 2 batches (for the design of batches and groups, see Supplementary Tables 1-2). In methods that allow for covariates (namely, *DEXSeq, DRIMSeq, satuRn*, and *BRIE2*), we explicitly modelled the batch effect. Fifth, in order to investigate false positive detections, we generated null simulated datasets, where no differential effect is introduced between groups. In all simulations, we performed basic filtering and analyzed genes/transcripts with at least 10 counts per group. For further details about the simulation design, see Supplementary Details.

**Table 1:** Single-cell real data analysis. Number of interesting genes present among the top 200 results, of each cell-type, returned by every method. Methods “ *sat*”, “ *DiffReg* “, and *DEX* refer to *satuRn, DEXSeq*, and *DifferentialRegulation*, respectively. “brain only” denotes the 97 genes which were only detected in human brain; “cerebral cortex” indicates the 180 genes which display high expression in the human cerebral cortex, compared to other regions of the brain; “excitatory neurons” represents the 30 genes associated to excitatory neurons; “PGP1” refers to the 3 genes linked to PGP1; “overall” gathers all 299 genes belonging to any of the previous lists.

| Genes | *sat_SC* | *DiffReg* | *DiffReg_Wald* | *BRIE2* | *eisaR* | *sat* | *DEX_TECs* | *DRIM* |
| --- | --- | --- | --- | --- | --- | --- | --- | --- |
| brain only | 8 | 3 | 3 | 0 | 0 | 0 | 0 | 0 |
| cerebral cortex | 8 | 7 | 7 | 0 | 0 | 1 | 0 | 0 |
| excitatory neurons | 3 | 2 | 1 | 1 | 0 | 0 | 0 | 0 |
| PGP1 | 0 | 2 | 2 | 0 | 1 | 0 | 0 | 0 |
| overall | 17 | 14 | 13 | 1 | 1 | 1 | 0 | 0 |

## 3 Results

### 3.1 Bulk simulation study

We benchmarked *DifferentialRegulation* against *eisaR*, which was developed to identify changes in splicing dynamics from bulk RNA-seq data, and various competitors that recently displayed good performance in detecting DAS from bulk RNA-seq data: *DRIMSeq, satuRn, SUPPA2*, and *DEXSeq* (Anders et al., 2012), which was used on both transcript estimated abundance (i.e., *DEXSeq_TECs*), and equivalence classes counts (i.e., *DEXSeq_ECs*). Figure 2 reports the receiving operating characteristic (ROC) curve for the main bulk simulations, and the number of false detections among the top-ranked transcripts, which is particularly relevant because top discoveries are usually selected for subsequent analyses by life scientists. In all simulated scenarios, *DifferentialRegulation, eisaR* and *SUPPA2* display good performance, in terms of sensitivity, specificity, and false positive detections among top-ranked transcripts. Of the three methods, *SUPPA2* is the most affected by DGE and DAS confounding effects, while *DifferentialRegulation*’s results are consistent also when increasing the strength of the DGE effect (Supplementary Figure 1). *DEXSeq_TECs* and *satuRn* also perform well, yet with lower statistical power. Notably, *DRIMSeq* and *DEXSeq_ECs* display a low TPR, because they fail to analyze several transcripts, and return multiple NA’s. These results are consistent with what was previously observed in Love et al. (2018), Tiberi and Robinson (2020), and Gilis et al. (2022). When introducing batch effects, although the performance of all approaches decreases, the relative ranking of methods remains stable (Supplementary Figure 2). Further-more, *DifferentialRegulation* displays robust results, and leads to fewer false positive detections among its top results than competitors. In addition, we investigated how overall transcript abundance affects performance, and stratified the results of our main simulations into lowly, medium, and highly abundant transcripts, corresponding to the first, second, and third tertile of abundance, respectively. In general, higher abundance corresponds to increased statistical power, and brings the performance of all methods closer; in all cases, the relative ranking of methods remains approximately stable (Supplementary Figures 3-4).

**Figure 2:**
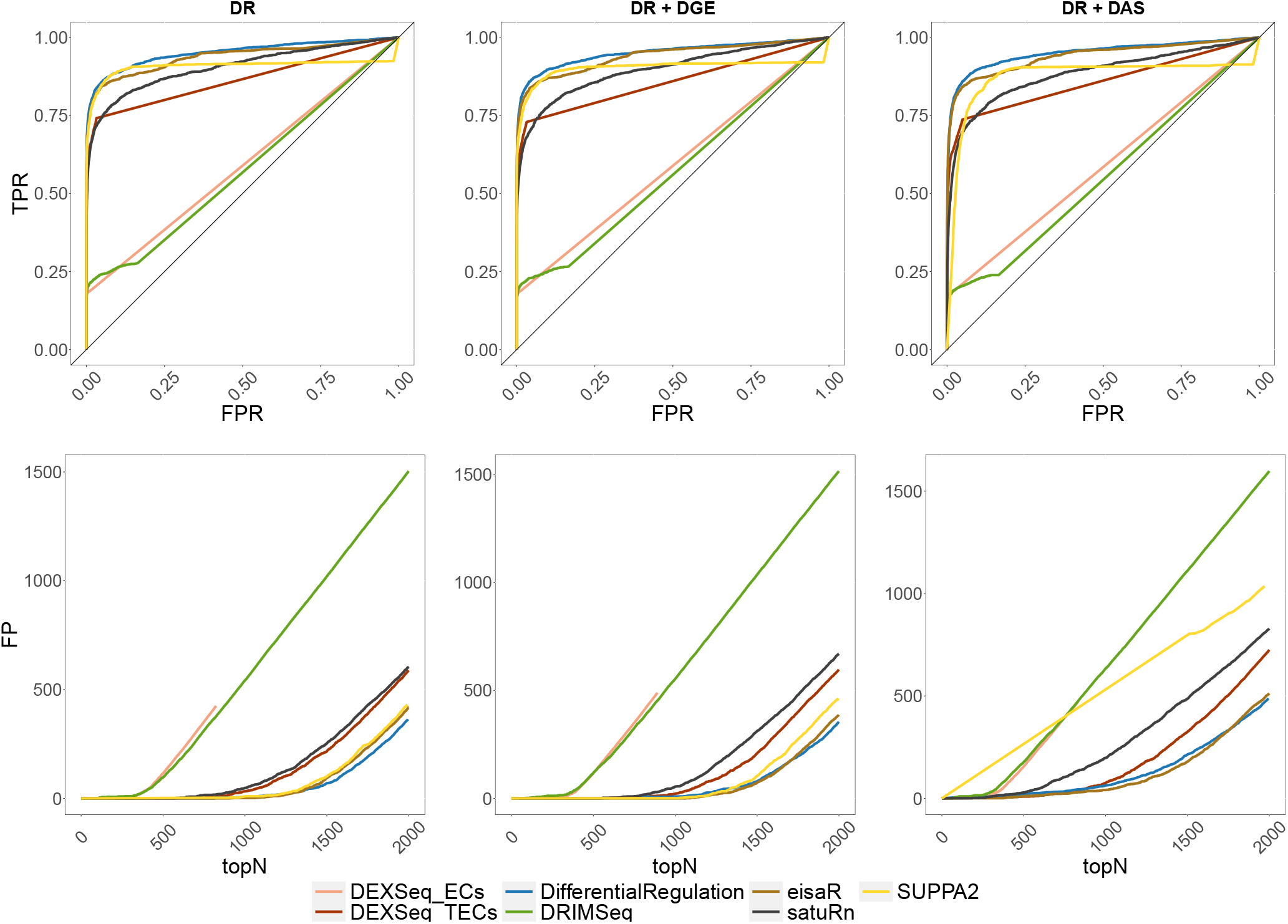
Results from the main bulk simulations. Top row: ROC curves; i.e., false positive rate (FPR) *vs*. true positive rate (TPR). Bottom row: false positive (FP) results among top detections (topN). Left panel (DR): simulation with differential regulation only; middle panel (DR + DGE): simulation with differential regulation and DGE (average fold-change of 3); right panel (DR + DAS): simulation with differential regulation and DAS.

**Figure 3:**
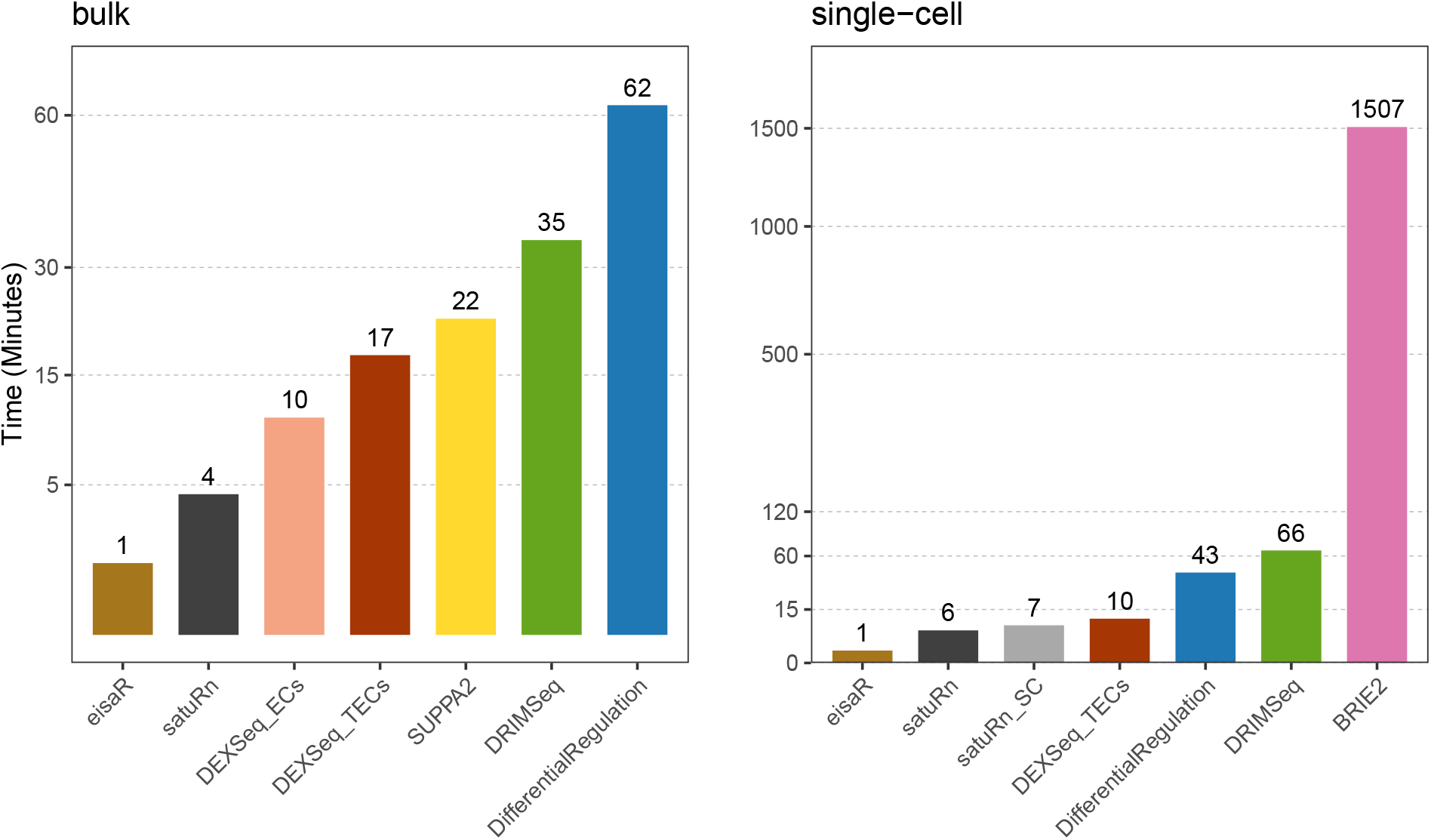
Average runtime (in minutes), of each method, across three main bulk (left panel; DR, DR + DGE, and DR + DAS, as in Figure 2), and the two single-cell main simulations (right panel; DR, and DR + DGE, as in Figure 4). All methods used 1 core, except *BRIE2*, which used 6 because the number of threads cannot be controlled by users.

In the null simulated dataset, all methods control well false positive detections: Supplementary Table 3 reports, for each method, the percentage of (raw) p-values below 0.1, 0.05 and 0.01. Additionally, *DifferentialRegulation*’s *p*, the estimated probability that group *B* is up-regulated (while 1 − *p* is the probability that *B* is down-regulated), is centred around 0.5, as one would expect when two groups are not differential (Supplementary Figure 5; left panel).

From a computational perspective, *DifferentialRegulation* is the most demanding method, which is unsurprising given the high cost of full MCMC algorithms involving latent states (Figure 3, left panel); nonetheless, the approach ran in about 1 hour on a single thread.

### 3.2 Single-cell simulation study

In the single-cell simulation, we benchmarked *DifferentialRegulation* against *BRIE2* and *satuRn_SC* (i.e., *satuRn* in its single-cell variant), and several approaches originally designed for bulk data: *eisaR, DEXSeq_TECs, DRIMSeq*, and *satuRn* (bulk variant). Except *BRIE2* and *satuRn_SC*, which use single-cell observations, all methods worked with pseudo-bulk counts (i.e., aggregated counts across cells in a cluster). Here, *DEXSeq_ECs* and *SUPPA2* were excluded because they could not be adapted to single-cell data: the former approach is bound to the output structure from *salmon*, which is a bulk pseudo-aligner, and the latter requires transcript-level abundances, while single-cell aligners (e.g., *alevin-fry*) return counts at the gene level. While *eisaR* was used on *s* and *u* reads, all remaining methods were run on *s, u* and *a* estimated counts, hence accounting for the uncertainty in ambiguous reads. Note that, however, only *DifferentialRegulation* accounts for the variability in reads mapping to multiple genes. In our main simulations, *DifferentialRegulation* displays good sensitivity and specificity, although its ROC curve is mainly below those of *eisaR* and *DEXSeq_TECs* (Figure 4). Nonetheless, our method has fewer false discoveries than competitors among top-ranked genes, particularly when introducing DGE as a nuisance effect (Supplementary Figure 6). Our approaches based on the posterior probability *p* (*DifferentialRegulation*), and on a Wald test (*DifferentialRegulation_Wald*) perform similarly, although the second one leads to (marginally) fewer false discoveries in top ranked genes (Figure 4). Results are consistent also when varying the fraction of cells with zero abundance (Supplementary Figure 7), and when dealing with batch effects (Supplementary Figure 8). In particular, while increasing zero abundance cells has little impact on the results (which is reasonable in pseudo-bulk methods), batch effects lead to a general deterioration in performance, particularly for *DRIMSeq, satuRn*, and *satuRn_SC*. Nonetheless, *DifferentialRegulation* results are robust, especially when considering top detections. As in the bulk simulation, we also stratified the results of our main simulations by overall gene expression, and found a consistent ranking of methods across abundance level; as expected, higher gene expression (i.e., more data) is associated with higher statistical power (Supplementary Figures 9-10). In the null simulation study, all methods (except *satuRn* and *satuRn_SC*) display a good control of false positives (Supplementary Table 4), and *DifferentialRegulation*’s *p* is again centred around 0.5, although with more variability compared to the bulk simulation (Supplementary Figure 5; right panel). Computationally, coherently with what previously observed, *eisaR, satuRn* and *DEXSeq_TECs* emerged as the fastest methods, while *DifferentialRegulation* required significantly more time, yet approximately 35 times less than the other Bayesian approach, *BRIE2* (Figure 3, right panel), despite *BRIE2* using 6 times more cores than any other method.

**Figure 4:**
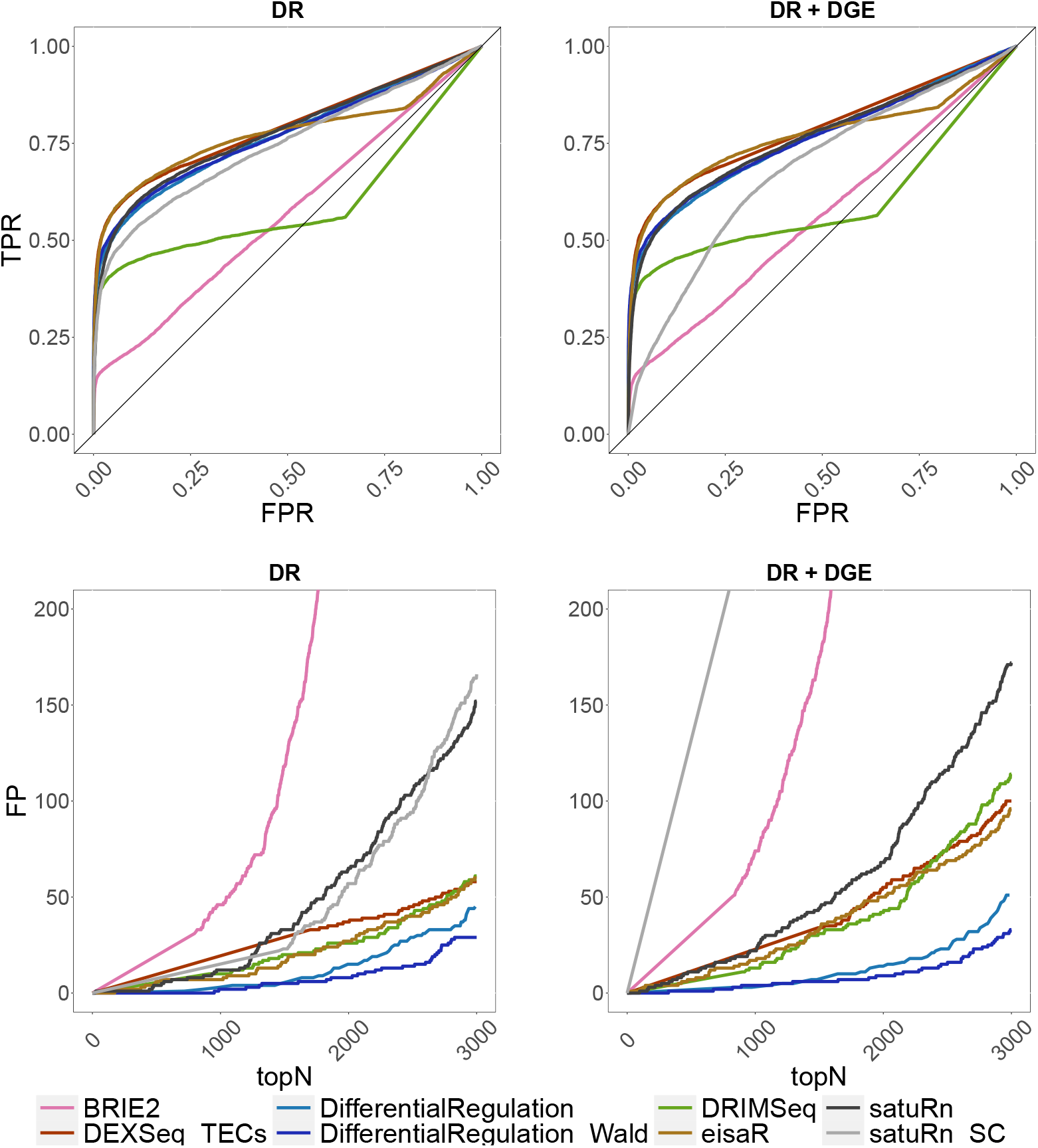
Results from the main single-cell simulations. Top row: ROC curves; i.e., false positive rate (FPR) *vs*. true positive rate (TPR). Bottom row: false positive (FP) results among top detections (topN). Left panel (DR): simulation with differential regulation only; right panel (DR + DGE): simulation with differential regulation and DGE (average fold-change of 3).

### 3.3 Real data application

To compare methods on a real dataset, we considered the scRNA-seq data from Velasco et al. (2019), containing a total of 21 brain organoids from the human cerebral cortex, which were grown *in vitro*, for up to 6 months. Here we only considered a subset of 6 brain organoids from the PGP1 stem cell line: 3 organoids were observed at three months of development, and 3 were collected at six months of development. Comparing these two groups of samples should highlight changes that happen during brain development. After filtering low quality cells, via the *scater* (McCarthy et al., 2017) R package, and lowly abundant genes, with less than 10 non-zero cells, we were left with a total of 35,972 genes and 25,556 cells. Using the cell-type annotation available from the original study (Velasco et al., 2019), we grouped cells in six cell types (Supplementary Table 5). We applied differential methods and discovered differences, for each cell type, across development time points. We then used The Human Protein Atlas website (Pontén et al., 2008) to generate the following lists of potentially interesting genes: 97 genes which have only been detected in the human brain; 180 genes that displayed significantly higher abundance in the human cerebral cortex (i.e., the area the organoids are derived from), compared to other regions of the brain; 30 genes associated to excitatory neurons, which play a key role in the development of the human brain cortex (Costa and Müller, 2015); 3 genes linked to the PGP1 cell line (i.e., the cell line used to generate the data). In absence of a ground truth, for each method, we investigated how often these genes appear in the top 200 ranked genes from each cell type. Overall, *satuRn_SC, DifferentialRegulation* and *DifferentialRegulation_Wald* top discoveries contain significantly more potentially interesting genes than competitors (Table 1). *DifferentialRegulation*’s results are coherent with the fact that, in the simulation studies, our approach, among its top ranked genes/transcripts, led to fewer false positives than other methods.

We also considered a bulk (paired-end) RNA-seq real dataset from pancreatic *β* cells in mouse embryos, collected during pancreas development (Osipovich et al., 2021). The dataset consists of 3 wild-type, and 3 Zfp800 knockout mice, where Zfp800 is a crucial protein for pancreas development. After removing lowly abundant transcripts (less than 10 counts in at least one group), we analyzed 54,904 transcripts, associated to a total of 16,347 distinct genes. Following a similar approach to the one outlined above, we built two lists of potentially interesting genes, searching for terms “mouse pancreas development” (110 genes) and “ZNF800” (3 genes) on The Human Protein Atlas. We then converted each transcript name to the name of the corresponding gene, and counted how many of each method’s top 1,000 results belong to these two lists. Compared to the single-cell real data, results are more homogeneous across methods (Supplementary Table 6); nonetheless, *DifferentialRegulation* identifies slightly more interesting genes than competitors. Finally, in both bulk and single-cell real data analyses, we visually checked convergence and mixing of the posterior chains for 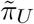 (i.e., the key parameter of inference) for the 20 most significant results (traceplots in Supplementary Figures 11-14). Additionally, we investigated how robust results are when running the algorithm multiple times. To this aim, we analyzed each real dataset twice, using distinct seeds for the random number generator (i.e., *set*.*seed* in R), and compared results: *Wald* test p-values across runs were highly coherent, with a Pearson correlation of 0.987 and 0.996 in the bulk and single-cell application, respectively.

## 4 Discussion

We have introduced *DifferentialRegulation*, a Bayesian hierarchical approach to discover differentially regulated genes and transcripts across conditions, by detecting changes in the relative abundance of unspliced reads, which indicate differences in the future mRNA production. Our method works with both bulk and single-cell RNA-seq data, and is based on two distinct models to adapt to the peculiar aspects of the data being analyzed. Similarly, the outputs of the two frameworks differ, and take advantage of the information that each data type provides: in bulk data, we target transcript-level changes (across all cells), while in single-cell data, we aim at cluster (e.g., cell-type) specific changes, yet at the gene level. Importantly, RNA-seq data is typically characterized by a high degree of quantification uncertainty: we account for it via a latent variable approach where reads are allocated to their gene/transcript of origin, and to the corresponding splice version.

Starting from real data as anchor data, we designed several benchmarks for bulk and single-cell RNA-seq data, and compared *DifferentialRegulation* to state-of-the-art tools. Our method displays good sensitivity and specificity, and shows fewer false discoveries than competitors among top ranked genes, that are usually chosen by biologists for further investigations. Additionally, our approach appears to be robust with respect to nuisance effects, such as differential gene expression, differential alternative splicing, and batch effects, and shows good performance even in lowly abundant genes/transcripts. We also performed two real data analyses, where our method can detect (among its top ranked genes) more potentially interesting genes than most alternative approaches.

We distributed *DifferentialRegulation*, open-access, as an R package via the Bioconductor project, which facilitates its integration with other bioinformatics tools and pipelines; furthermore, we provided an example usage vignette, and plotting functions that simplify the visualization of results.

Finally, we would like to acknowledge some limitations of our framework. First, our method is amongst the most computationally demanding tools we tested, although clever coding techniques (such as under-sampling, and C++ coding) enabled us to run our approach in our benchmarks in approximately 40-60 minutes using a single core. Furthermore, note that, our single-cell approach can also benefit from parallel coding, which can be particularly useful in large datasets. Second, covariates, such as batch effects, are not modelled; nonetheless, such nuisance effects usually affect overall gene abundance, while our framework focuses on relative abundance, and (as shown) is robust to DGE changes; therefore, such covariates are unlikely to impact results.

## Availability

*DifferentialRegulation* is freely available as a Bioconductor R package at https://bioconductor.org/packages/DifferentialRegulation. The scripts used to run all analyses are available on GitHub (https://github.com/SimoneTiberi/DifferentialRegulation_manuscript, release v2). R scripts were run with R version 4.3.0, and Bioconductor packages from release 3.17. Raw data (fastq files) for the sample used to seed the bulk simulation are available from https://www.ebi.ac.uk/ena/browser/view/SRR493366. The dataset used as anchor for the single-cell simulation can be downloaded from Gene Expression Omnibus (accession number GSE107585). The bulk data used in the real data analysis is available at the Gene Expression Omnibus (accession number GSE129519). The single-cell data used in the real data analysis is available at ArrayExpress (accession number E-MTAB-9538).

## Acknowledgements

We acknowledge Amanda Kedaigle, the Robinson lab, and our reviewers for their feedback and suggestions. This work was supported by Swiss National Science Foundation grants 310030_175841 and 310030_204869 to MDR. MDR acknowledges support from the University Research Priority Program Evolution in Action at the University of Zurich. RP acknowledges NIH award R01HG009937 and NSF awards CCF-1750472 and CNS-1763680.

## Author contributions

JM, PC and CS performed some of the analyses. DH, HS and RP contributed to the simulation studies. AA contributed to the method development. MDR contributed to designing the study. ST designed the study, implemented the method, performed the revisions and most of the analyses, and wrote the manuscript. All authors contributed to editing the manuscript, and approve the article.

