## Supplementary material for "*DifferentialRegulation*: a Bayesian hierarchical approach to identify differentially regulated genes"

[1] *Department of Statistical Sciences, University of Bologna, Bologna, Italy.*
[2] *Department of Molecular Life Sciences and SIB Swiss Institute of Bioinformatics, University of Zurich, Zurich, Switzerland.*
[3] *Computational Biology Platform, Friedrich Miescher Institute for Biomedical Research and SIB Swiss Institute of Bioinformatics, Basel, Switzerland*
[4] *Department of Computer Science, University of Maryland, MD, USA*
[5] *Department of Computer Science, Princeton University, NJ, USA*
[6] *Research Unit of Applied Statistics, TU Wien, Vienna, Austria*
[7] *Harvard-MIT Center for Regulatory Science, Harvard Medical School, Boston, MA, USA*
[8] *Department of Cell Biology and Molecular Genetics, University of Maryland, MD, USA*
[9] *Center for Bioinformatics and Computational Biology, University of Maryland, MD, USA*

#### 1 Supplementary Details

##### 1.1 Ambiguous reads in single-cell RNA-seq data

When using single-cell RNA-seq data, we only employ a latent variable approach for multi-gene mapping reads, while $a$ reads are not allocated, and are treated separately from $s$ and $u$. This is because, in order to allocate $a$ reads, we need an estimate of the probability that an ambiguous read is spliced (or unspliced), which cannot be accurately computed, because it depends on various unknown factors.

In fact, existing single-cell analysis tools either assign all ambiguous reads as spliced, for example, *velocyto* (La Manno et al., 2018), or acknowledge the difficulty and quantify the result of the ambiguous reads as a stand-alone count matrix (i.e., $a$), and leave the task of solving the ambiguity to the users, such as *alevin* (Srivastava et al., 2019) and *alevin-fry* (He et al., 2022). The difficulty of ambiguous read allocation comes from both biological and computational aspects. First, $s$ and $u$ mRNA share large genomic regions (e.g., exons), while scRNA-seq reads are characterized by short fragments, which are often compatible with both splice variants of genes. Second, most ambiguous reads can be allocated to spliced and unspliced genes equally well, with no decisive evidence for disambiguating those reads. Third, while read coverage is approximately uniform in bulk RNA-seq reads (i.e., the location of reads in a transcript can be assumed to be roughly random, hence uniform, across the transcript), this is not the case for single-cell protocols. At present, we still lack accurate methods that model the underlying mechanism of ambiguous reads, and address the respective uncertainty in a probabilistic way.

##### 1.2 MCMC scheme

We alternately sample parameters and latent states from their conditional distributions, following a Metropolis-within-Gibbs sampling scheme. In particular, we sample:

- hyper-parameters $\delta|\pi$ from a Metropolis sampler, with an adaptive random walk proposal;
- parameters $\rho|X$ from Dirichlet Gibbs samplers;
- hierarchical parameters $\pi|X, \delta$ from Dirichlet Gibbs samplers;
- latent states $X|Z, \pi$ from multinomial Gibbs samplers.

Below, we describe the prior specifications, and the details of each sampling scheme.

##### 1.2.1 $\delta|\pi$ Metropolis sampling

For convenience, the $\delta$ parameters are sampled after applying a logarithmic transformation. The posterior distribution of $log(\delta)$ is defined as: $p(log(\delta)|\pi) \propto p(\pi|log(\delta))p(log(\delta))$, where $p(log(\delta))$ is the informative prior for $log(\delta)$ (defined below). Instead, $p(\pi|log(\delta))$, in the bulk case, is the product of the $T$ beta densities defined in (2.3) (main paper), while, in the single-cell model, is the product of the $G$ Dirichlet densities defined in (2.6) (main paper).

We use an empirical Bayes approach to formulate an informative prior for the hyper-parameters. In particular, we randomly select 1,000 genes/transcripts, and, for each one of them, we use *DRIMSeq* (Nowicka and Robinson, 2016) to obtain an initial estimate of the following parameters: $log(\delta_+), log(\delta_S)$, and (in the single-cell model) $log(\delta_U)$. For each parameter, we compute the mean and standard-deviation of its 1,000 estimated values (1 per selected gene): $\mu_+$ and $\sigma_+$ for $log(\delta_+)$, $\mu_S$ and $\sigma_S$ for $log(\delta_S)$, and (in the single-cell model) $\mu_U$ and $\sigma_U$ for $log(\delta_U)$.

These estimates are used to formulate the following informative prior distributions for the hyper-parameters: $log(\delta_+) \sim \mathcal{N}\left(\mu_+, \sigma_+^2\right)$, $log(\delta_S) \sim \mathcal{N}\left(\mu_S, \sigma_S^2\right)$, and (in the single-cell model) $log(\delta_U) \sim \mathcal{N}\left(\mu_U, \sigma_U^2\right)$.

Hyper-parameters are sampled, in the logarithmic scale, from a Metropolis sampler; in particular, we sample parameters $(log(\delta_S), log(\delta_U), log(\delta_A))$, (or simply $(log(\delta_S), log(\delta_U))$ in the bulk model), and, to obtain their prior distributions, apply the change of variable via the Jacobian transformation (Murphy, 2012).

Values are proposed based on an adaptive random walk (Haario et al., 2001). In the initial 200 iterations, we propose values from a normal distribution independently for each hyper-parameter. After 200 iterations, a covariance matrix of the posterior chains is computed, and hyper-parameters are proposed from a multivariate normal distribution, with this covariance matrix. The covariance matrix is then re-updated at 300, 400, 500 iterations, and at the burn-in (which is set to be at least 500 iterations).

##### 1.2.2 $\rho|X$ Gibbs sampling

Below, we express densities in terms of $Y$, which is directly derived from $X$: in the bulk model, for each transcript $t$, and sample $i$, $Y_i^{(t)}$ is obtained as $Y_i^{(t)} = X_{Si}^{(t)} + X_{Ui}^{(t)}$. Similarly, in the single-cell model, for gene $g$ and sample $i$, we have that $Y_i^{(g)} = X_{Si}^{(g)} + X_{Ui}^{(g)} + X_{Ai}^{(g)}$.

We assume a weakly informative Dirichlet prior distribution for $\rho$,

$$\rho_i \sim Dir\left(1, \ldots, 1\right), \text{ for } i = 1, \ldots, N. \tag{1}$$

The posterior distribution of $\rho_i$ can be obtained as: $p(\rho_i|Y_i) \propto p(Y_i|\rho_i)p(\rho_i)$, where $p(Y_i|\rho_i)$ is the density of the multinomial distribution, reported in the main paper in formula (2.1) for the bulk model, and (2.4) for the single-cell model. Given the conjugacy of the Dirichlet and multinomial distributions, the posterior distribution is again Dirichlet distributed, and, in the bulk model, it is equal to:

$$\rho_i|Y_i \sim Dir\left(Y_i^{(1)} + 1, \ldots, Y_i^{(T)} + 1\right) \text{ for } i = 1, \ldots, N. \tag{2}$$

In the single-cell model, $G$ (the number of genes) replaces $T$ (the number of transcripts).

###### 1.2.3 $\pi | X, \delta$ Gibbs sampling

In the bulk model, for each sample $i$ and transcript $t$, the posterior distribution of $\pi_{Si}^{(t)}$ can be obtained as: $p\left(\pi_{Si}^{(t)} \middle| X_{Si}^{(t)}, \delta^{(t)}\right) \propto p\left(X_{Si}^{(t)} \middle| \pi_{Si}^{(t)}\right) p\left(\pi_{Si}^{(t)} \middle| \delta^{(t)}\right)$, where $p\left(X_{Si}^{(t)} \middle| \pi_{Si}^{(t)}\right)$ is the density of the Binomial distribution in (2.2) (main paper), and $p\left(\pi_{Si}^{(t)} \middle| \delta^{(t)}\right)$ is the conjugate Beta hierarchical prior distribution in (2.3) (main paper). These choices lead to the following Beta posterior distribution for $\pi_{Si}^{(t)}$:

$$\pi_{Si}^{(t)} \Big| X_i^{(t)}, \delta^{(t)} \sim Beta\left(X_{Si}^{(t)} + \delta_S^{(t)}, X_{Ui}^{(t)} + \delta_U^{(t)}\right), \text{ for } t = 1, \dots T, i = 1, \dots N. \tag{3}$$

Similarly, in the single-cell model, for each sample $i$ and gene $g$, the posterior distribution of $\pi_i^{(g)}$ can be calculated as: $p\left(\pi_i^{(g)} \middle| X_i^{(g)}, \delta^{(g)}\right) \propto p\left(X_i^{(g)} \middle| \pi_i^{(g)}\right) p\left(\pi_i^{(g)} \middle| \delta^{(g)}\right)$, where $p\left(X_i^{(g)} \middle| \pi_i^{(g)}\right)$ is the density of the multinomial distribution in (2.5) (main paper), and $p\left(\pi_i^{(g)} \middle| \delta^{(g)}\right)$ is the conjugate Dirichlet hierarchical prior distribution in (2.6) (main paper). These results lead to the following Dirichlet posterior distribution for $\pi_i^{(g)}$:

$$\pi_i^{(g)} \Big| X_i^{(g)}, \delta^{(g)} \sim Dir\left(X_{Si}^{(g)} + \delta_S^{(g)}, X_{Ui}^{(g)} + \delta_U^{(g)}, X_{Ai}^{(g)} + \delta_A^{(g)}\right), \text{ for } g = 1, \dots G, i = 1, \dots, N. \tag{4}$$

###### 1.2.4 $X | Z, \pi$ Gibbs sampling

Assume that a total of $J$ equivalence classes (EC) are available. Each EC is defined by two elements, which for the $j$-th EC are $W_j$ and $V_j$: the former contains the list of gene(s)/transcript(s) compatible with the $j$-th equivalence class, while the latter indicates the splice status of the corresponding gene(s)/transcript(s) (i.e., $s$ or $u$ for bulk data, and $s$, $u$ or $a$ for single-cell data). Each EC is also associated to a count, $z_j$ for the $j$-th class, indicating the number of (bulk or single-cell) RNA-seq reads compatible with it.

For each EC, we aim to allocate its counts to the gene(s)/transcript(s), and respective splice status, compatible with it.

For simplicity, below we consider the bulk model; the single-cell model is a natural extension of it. We define $X_{Sij}^{(t)}$ and $X_{Uij}^{(t)}$ as the contribution (i.e., counts) of the $j$-th EC to the spliced and unspliced versions of transcript $t$ in sample $i$.

Assume that the $j$-th EC is compatible with $n_j$ transcripts. The counts of the $j$-th EC from the $i$-th sample are allocated according to the following multinomial distribution:

$$\left(X_{v_1 i}^{(w_1)}, \dots, X_{v_{n_j} i}^{(w_{n_j})}\right) \Bigg| \pi_i, Z \sim \mathcal{MN}\left(\pi_{ij}, z_j\right), \text{ for } i = 1, \dots, N, j = 1, \dots, J, \tag{5}$$

where $\pi_{ij} \propto \left(\rho_i^{(w_1)} \times \tilde{\pi}_{v_1 i}^{(w_1)}, \dots, \rho_i^{(w_{n_j})} \times \tilde{\pi}_{v_{n_j} i}^{(w_{n_j})}\right)$, $W_j = \left(w_1, \dots, w_{n_j}\right)$, and $V_j = \left(v_1, \dots, v_{n_j}\right)$, with $w_1, \dots, w_{n_j} \in \{1, \dots, T\}$, and $v_1, \dots, v_{n_j} \in \{S, U\}$.

For every sample, $i = 1, \dots, N$, once the counts of all $J$ ECs have been allocated, we can recover the overall $s$ and $u$ abundance of each transcript, $t = 1, \dots, T$, by adding elements across ECs: $X_{Si}^{(t)} = \sum_{j=1}^J X_{Sij}^{(t)}$ and $X_{Ui}^{(t)} = \sum_{j=1}^J X_{Uij}^{(t)}$; then, the overall transcript-level counts can be retrieved as $Y_i^{(t)} = X_{Si}^{(t)} + X_{Ui}^{(t)}$.

In order to accelerate the runtime of our method, latent states are only sampled every 10 iterations (i.e., undersampling scheme).

##### 1.3 Wald-type test for the single-cell model

Given two groups of samples, $A$ and $B$, we aim to test if parameter $\pi_S$, $\pi_U$, and $\pi_A$ change between groups. We consider the difference of these parameters across groups as: $\omega = (\omega_S, \omega_U)$, with $\omega_S = {}_A\pi_S - {}_B\pi_S$, and

$\omega_U = {}_A\pi_U - {}_B\pi_U$. Under the null hypothesis, $\pi_S$ and $\pi_U$ (and consequently $\pi_A$) do not change across groups; therefore we test the following system of hypotheses: $H_0 : \omega_S = \omega_U = 0$; $H_1$ : otherwise.

We approximate the posterior distribution of $\omega$ with a bivariate normal (Gelman et al., 1995): $\omega \dot{\sim} \mathcal{N}(\hat{\omega}, \hat{\Sigma}_\omega)$, with parameters $\hat{\omega}$ and $\hat{\Sigma}_\omega$ inferred from the posterior chains. Given this normal approximation, we use a bivariate Wald test (Li et al., 1991): $\hat{\omega}\hat{\Sigma}_\omega^{-1}\hat{\omega}^T \dot{\sim} \chi^2_2$.

##### 1.4 Simulation details

In the main paper (Section 2.5), we illustrate how, in both bulk and single-cell simulations, starting from a real dataset as anchor data, we obtain the counts for each transcript (bulk data) or gene (single-cell data). Then, we artificially introduce a differential effect in a randomly selected subset of the genes/transcripts. Finally, starting from these counts, in order to introduce mapping uncertainty, we use read-level simulators (*RSEM* (Li and Dewey, 2011) for bulk data and *minnow* (Sarkar et al., 2019) for single-cell data) to simulate read-level data. Below, we explain how the differential effects are introduced.

To generate differentially regulated (DR) genes/transcripts, we first define a list of genes/transcripts that are eligible for being chosen as DR. In particular, a gene/transcript is eligible when, if it was selected as DR, it would lead to a difference, between groups $A$ and $B$, in the (group-level) relative abundance of unspliced reads, $\bar{\pi}_U$, of at least 0.2 (i.e., ${}_B\bar{\pi}_U - {}_A\bar{\pi}_U \geq 0.2$). Since DR is introduced by inverting spliced and unspliced reads in one group, this condition can be easily checked, in the data where DR has not been introduced, as: ${}_B\bar{\pi}_S - {}_A\bar{\pi}_U \geq 0.2$. This condition prevents us from simulating negligible differential effects (i.e., ${}_B\bar{\pi}_U - {}_A\bar{\pi}_U < 0.2$), which would be uninteresting. Then, from the list of eligible genes/transcripts, we randomly select 2,000 transcripts (bulk simulation), or 20% of the genes (single-cell simulation). For each one of the selected genes/transcripts, we randomly select the group ($A$ or $B$) where we will alter count data; this ensures that, on average, half of the modifications will affect group $A$, and half group $B$. Finally, for each gene, in each sample belonging to the selected group, we invert their spliced ($s$) and unspliced ($u$) counts. This introduces a difference in the relative abundance of $s$ and $u$ reads between groups.

In our DGE simulation, additionally to DR genes/transcripts (see above for DR simulation design), we also introduce a nuisance DGE effect. To this aim, we randomly sample 2,000 (bulk simulation) or 20% of the genes (single-cell data). For each gene/transcript, we randomly sample a fold change $FC$ between conditions, and a group to alter: the gene counts in the selected group are then multiplied by $FC$ (and rounded to the closest integer). As for the $DR$ effect, randomly sampling the group ensures that, on average, changes will equally affect the two groups. In the simulations with average fold change 3, 6 and 9, $FC$ is sampled as $FC = 2 + Z$, $FC = 5 + Z$ and $FC = 8 + Z$, respectively, with $Z \sim exp(\mu = 1)$, where $exp(\mu)$ is the exponential random variable with mean $\mu$. This generates random fold changes with variance 1, and mean 3, 6 or 9.

In our DAS simulation (bulk data only), after simulating DR transcripts (details above), we generate a nuisance DAS effect. To do that, we first randomly sample 2,000 genes; in order to introduce DAS, for each selected gene, we randomly sample a group: in the selected group, we invert the abundance (i.e., counts) of the two most abundant transcripts in the gene. This is coherent with the design of previous simulation studies involving DAS (Tiberi and Robinson, 2020; Soneson et al., 2016). Note that DAS is only introduced in bulk simulations, because its simulation requires transcript-level resolution, which is not available in scRNA-seq protocols.

In the batch simulation study, we generate two DR effects: one between groups (of interest), and one between batches (nuisance). First, we specify a design for batches: see Supplementary Tables 1 and 2 for the designs of the bulk and single-cell data, respectively. Then, we generate a DR effect between groups (see details above), and a second DR effect between batches. Clearly, the two DR effects are based on distinct sets of randomly selected genes/transcripts. In each simulation, we simulate the same number of DR genes/transcripts between groups (difference of interest) and between batches (nuisance effect): 2,000 transcripts for the bulk data, and 20% of the genes for the single-cell data.

In the dropout simulation study (single-cell data only), we generate three simulations where, for each gene in every cell type, we only allow for 1, 5 and 10% of the cells to have non-zero abundance. In particular, for each gene and cell type, we add the abundance of $s$ and $u$ reads across all cells of the specific cell type; then, we randomly allocate these $s$ (and $u$) overall counts across a randomly selected fraction (1, 5 or 10%) of genes. In this way the overall $s$ and $u$ abundances, of each gene in every cell type, are unchanged. Finally, starting from this dataset, we introduce a DR effect between groups, as in the main DR simulation (details above), by inverting the abundance of $s$ and $u$ reads in 20% of the genes (selected at random).

The null simulation study is trivially obtained by simulating no differential effects of any kind between groups.

Finally, note that, although *BRIE2* can account for the sample-to-sample variability, we found that including sample information led to a major decrease in power (Supplementary Figure 15); therefore, in all results, we considered the results obtained by fitting *BRIE2* with group as the only covariate in the model.

### 2 Supplementary Tables

| sample | group | batch |
|---|---|---|
| sample 1 | group A | batch 1 |
| sample 2 | group A | batch 1 |
| sample 3 | group A | batch 2 |
| sample 4 | group B | batch 1 |
| sample 5 | group B | batch 2 |
| sample 6 | group B | batch 2 |

**Supplementary Table 1:** Design of the simulation study with batch effects; bulk data.

| sample | group | batch |
|---|---|---|
| sample 1 | group A | batch 1 |
| sample 2 | group A | batch 2 |
| sample 3 | group B | batch 1 |
| sample 4 | group B | batch 2 |

**Supplementary Table 2:** Design of the simulation study with batch effects; single-cell data.

| P-value threshold | 0.1 | 0.05 | 0.01 |
|---|---|---|---|
| *DEXSeq_ TECs* | 0.1 | 0.1 | 0.1 |
| *DEXSeq_ ECs* | 0.0 | 0.0 | 0.0 |
| *DRIMSeq* | 0.9 | 0.4 | 0.1 |
| *DifferentialRegulation* | 0.1 | 0.0 | 0.0 |
| *satuRn* | 5.3 | 2.2 | 0.4 |
| *SUPPA2* | 1.7 | 0.5 | 0.0 |
| *eisaR* | 1.2 | 0.4 | 0.0 |

**Supplementary Table 3:** Percentage of false positive results in the null simulation study (bulk data); i.e., raw p-values below thresholds 0.1, 0.05, and 0.01. For *DifferentialRegulation*, we considered a false positive detection when $p$, the probability of up-regulation, was close to 0 or 1: $p < 0.05$ or $p > 0.95$ (for threshold 0.1), $p < 0.025$ or $p > 0.975$ (for threshold 0.05), and $p < 0.005$ or $p > 0.995$ (for threshold 0.01).

| P-value threshold | 0.1 | 0.05 | 0.01 |
|---|---|---|---|
| *BRIE2* | 4.0 | 2.5 | 0.9 |
| *DEXSeq_ TECs* | 0.1 | 0.1 | 0.0 |
| *DifferentialRegulation* | 3.5 | 1.7 | 0.4 |
| *DifferentialRegulation_ Wald* | 1.5 | 0.8 | 0.3 |
| *DRIMSeq* | 1.3 | 0.6 | 0.2 |
| *eisaR* | 4.9 | 2.1 | 0.3 |
| *satuRn* | 34.4 | 23.2 | 9.2 |
| *satuRn_ SC* | 21.1 | 12.5 | 4.6 |

**Supplementary Table 4:** Percentage of false positive results in the null simulation study (single-cell data); i.e., raw p-values below thresholds 0.1, 0.05, and 0.01. For *DifferentialRegulation*, we considered a false positive detection when $p$, the probability of up-regulation, was close to 0 or 1: $p < 0.05$ or $p > 0.95$ (for threshold 0.1), $p < 0.025$ or $p > 0.975$ (for threshold 0.05), and $p < 0.005$ or $p > 0.995$ (for threshold 0.01).

| cell type | number of cells |
|---|---|
| oRG | 6776 |
| CPNs | 6361 |
| Immature CPNs | 5983 |
| Immature PNs | 2954 |
| Cycling | 2389 |
| RG | 1093 |

**Supplementary Table 5:** Number of cells available, across all six brain organoids, for each cell type, in the real data analysis study.

| Genes | *DiffReg* | *DEXSeq_ TECs* | *eisaR* | *SUPPA2* | *DRIMSeq* | *DEXSeq_ ECs* | *satuRn* |
|---|---|---|---|---|---|---|---|
| mouse pancreas development | 12 | 12 | 11 | 11 | 10 | 9 | 6 |
| ZNF800 | 1 | 0 | 0 | 0 | 0 | 0 | 0 |
| overall | 13 | 12 | 11 | 11 | 10 | 9 | 6 |

**Supplementary Table 6:** Bulk real data analysis. Number of interesting genes present among the top 1,000 results returned by every method. "*DiffReg*" refers to *DifferentialRegulation*; "mouse pancreas development" and "ZNF800" indicate the 110 genes associated to the term "mouse pancreas development", and the 3 genes associated to "Znf800", respectively; "overall" gathers all 112 genes from the previous two lists.

### 3 Supplementary Figures

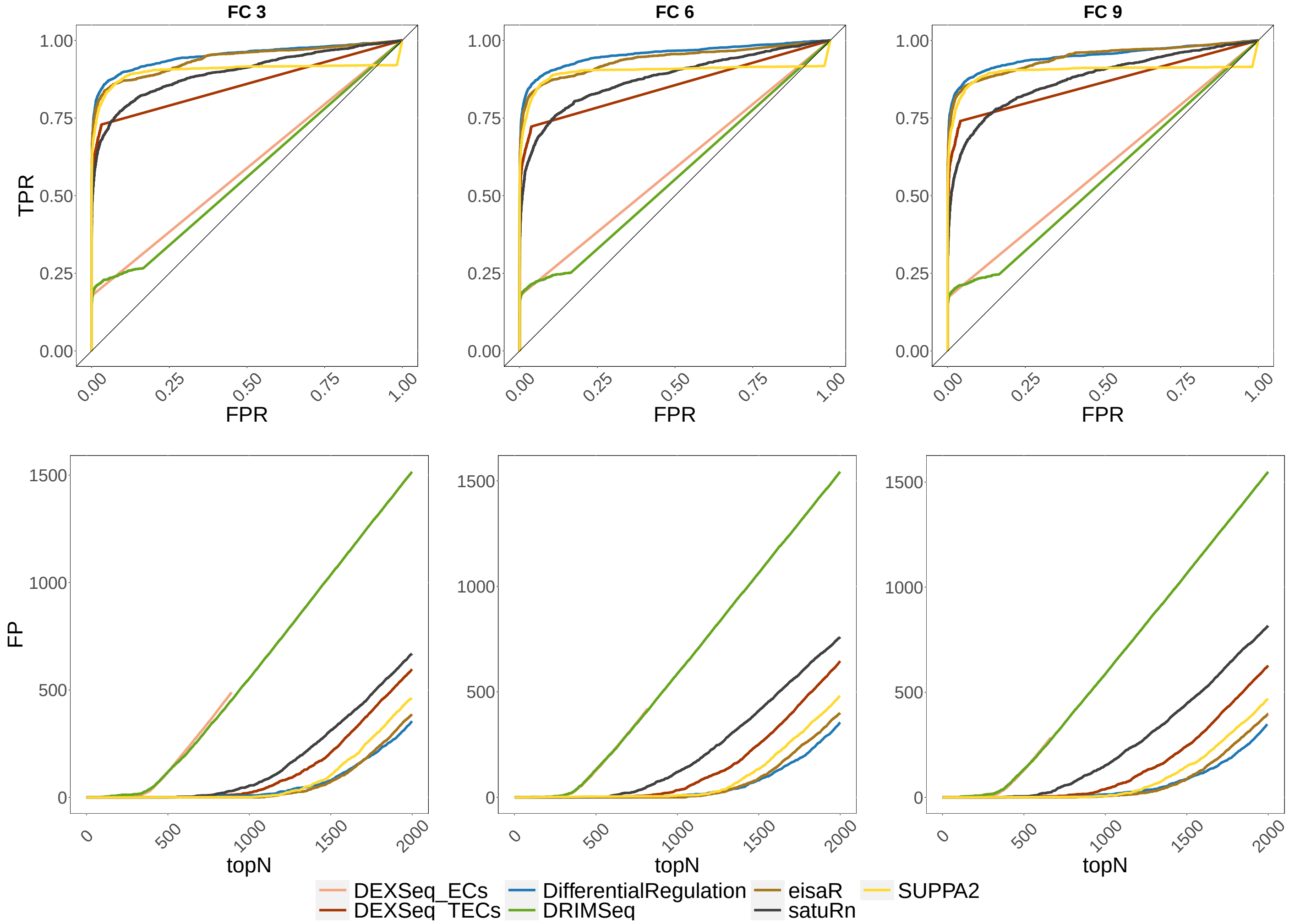


**Supplementary Figure 1:** Bulk simulation stud with DR + DGE. Top row: ROC curves; i.e., false positive rate (FPR) *vs.* true positive rate (TPR). Bottom row: false positive (FP) results among top detections (topN). All simulations have differentially regulated and differentially expressed genes, with average fold-change (FC) of 3 (left panel), 6 (middle panel) or 9 (right panel).

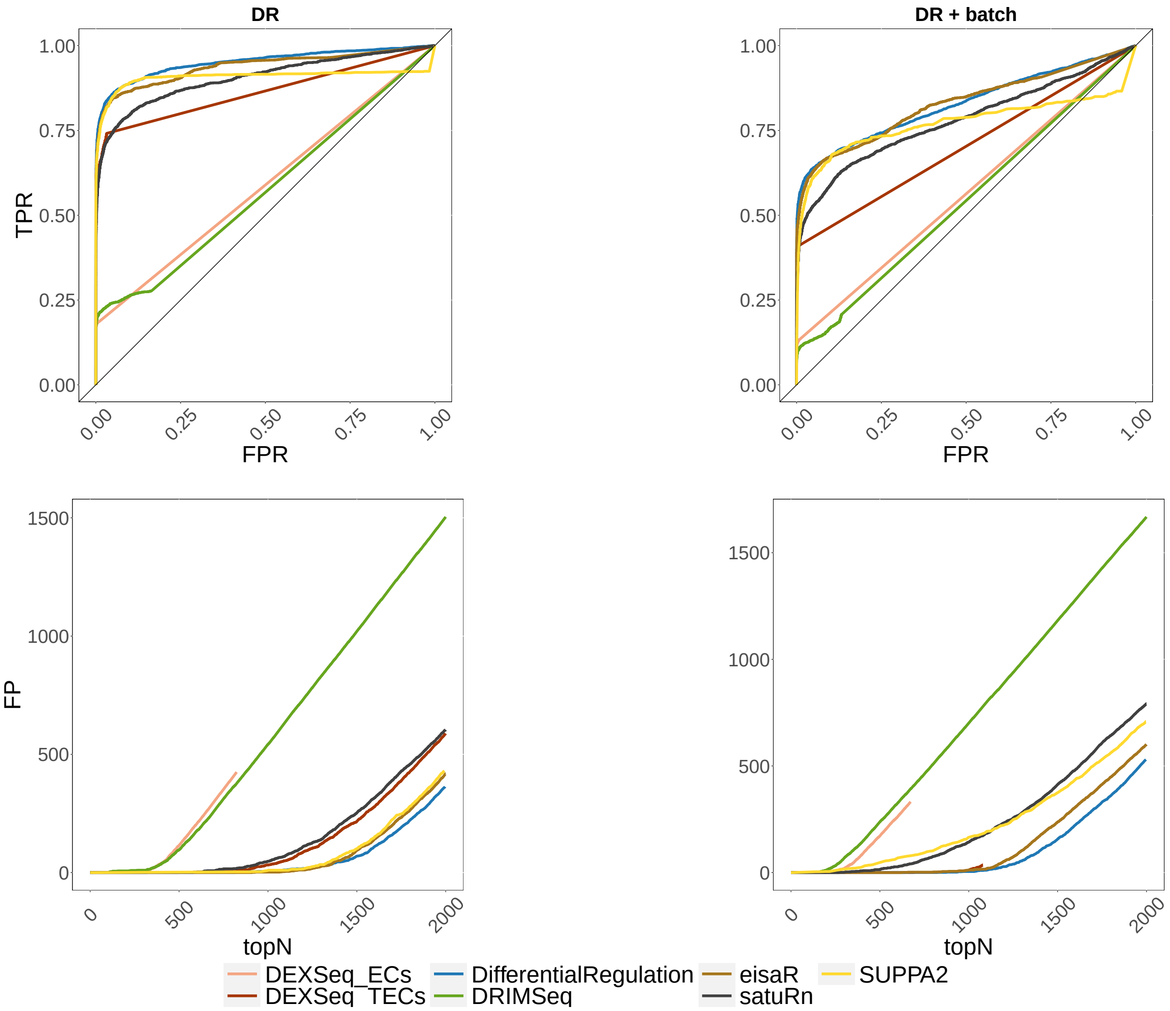


**Supplementary Figure 2:** Results from the bulk simulations, with and without batch effects. Top row: ROC curves; i.e., false positive rate (FPR) *vs.* true positive rate (TPR). Bottom row: false positive (FP) results among top detections (topN). Left panel (DR): simulation with differential regulation only; right panel (DR + batch): simulation with differential regulation and batch effect.

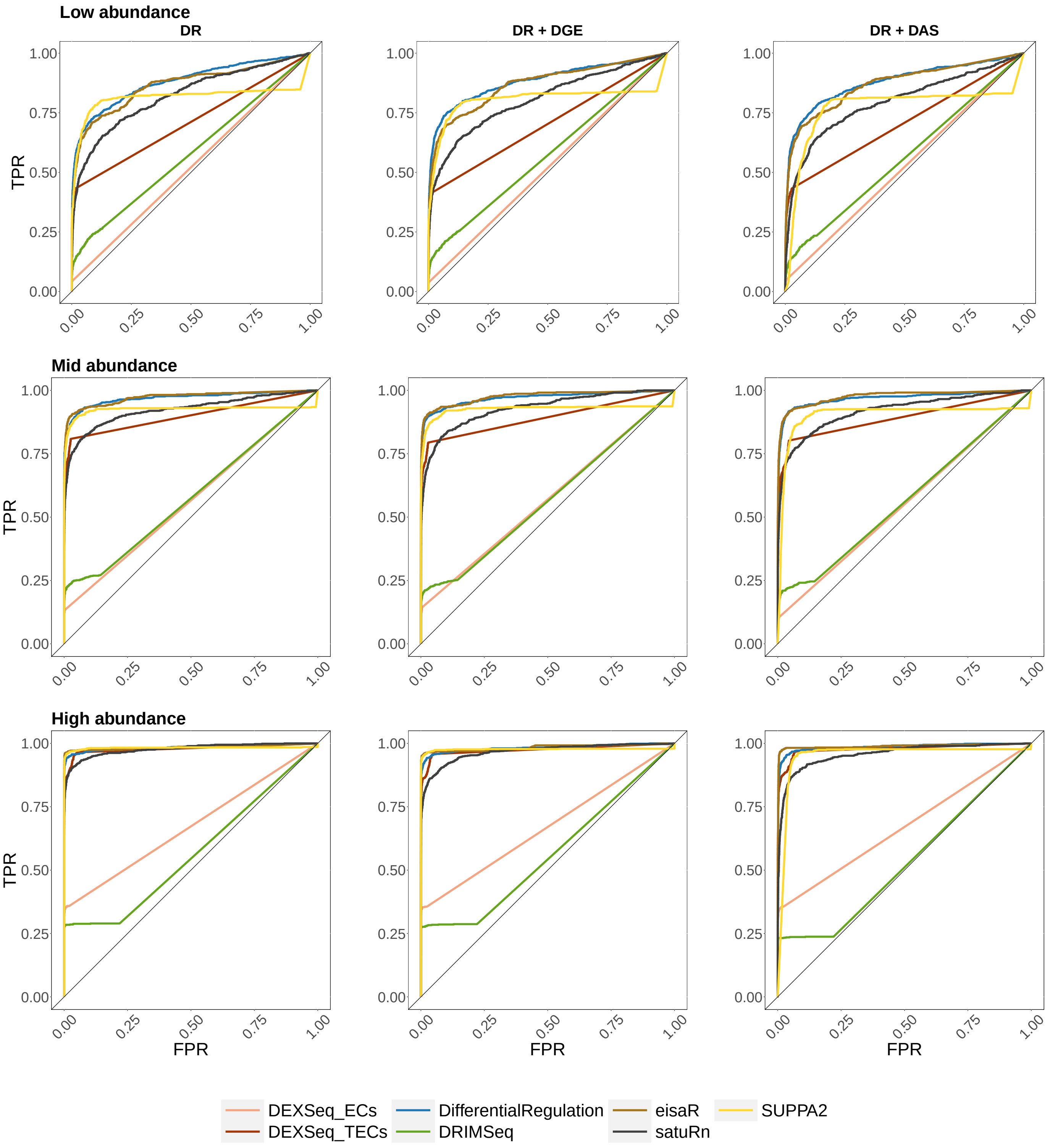


**Supplementary Figure 3:** ROC curves for the main bulk simulations, stratified by overall transcript abundance: false positive rate (FPR) *vs.* true positive rate (TPR). Top row: low abundance (bottom 33% of abundant transcripts); middle row: medium abundance (middle 33% of abundance); bottom row: high abundance (top 33% of abundant transcripts). Left column: DR only simulation; middle column: DR and DGE simulation (average fold-change of 3); right column: DR and DAS simulation.

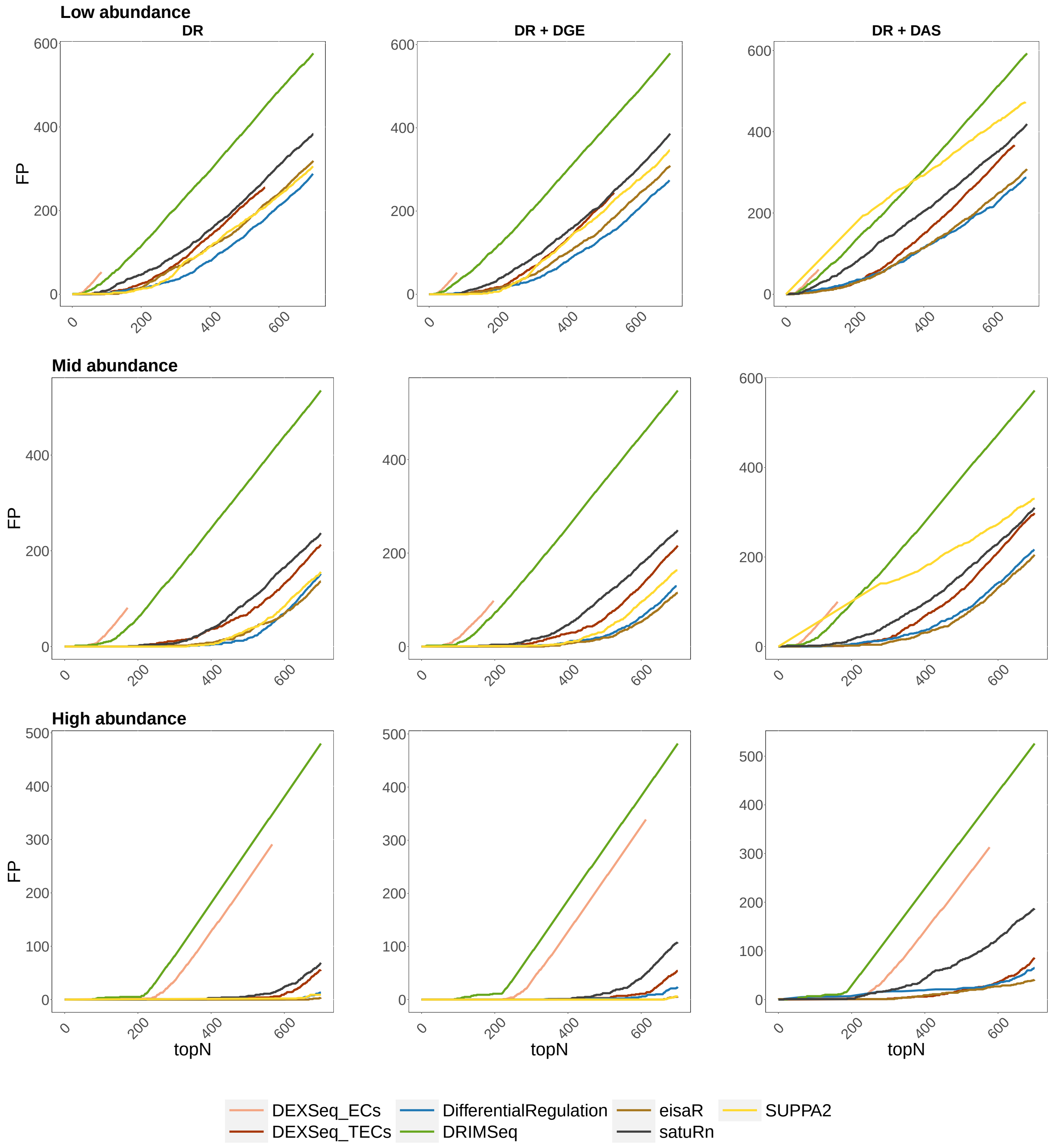


**Supplementary Figure 4:** False positive (FP) results among top detections (topN) for the main bulk simulations, stratified by overall transcript abundance: false positive rate (FPR) *vs.* true positive rate (TPR). Top row: low abundance (bottom 33% of abundant transcripts); middle row: medium abundance (middle 33% of abundance); bottom row: high abundance (top 33% of abundant transcripts). Left column: DR only simulation; middle column: DR and DGE simulation (average fold-change of 3); right column: DR and DAS simulation.

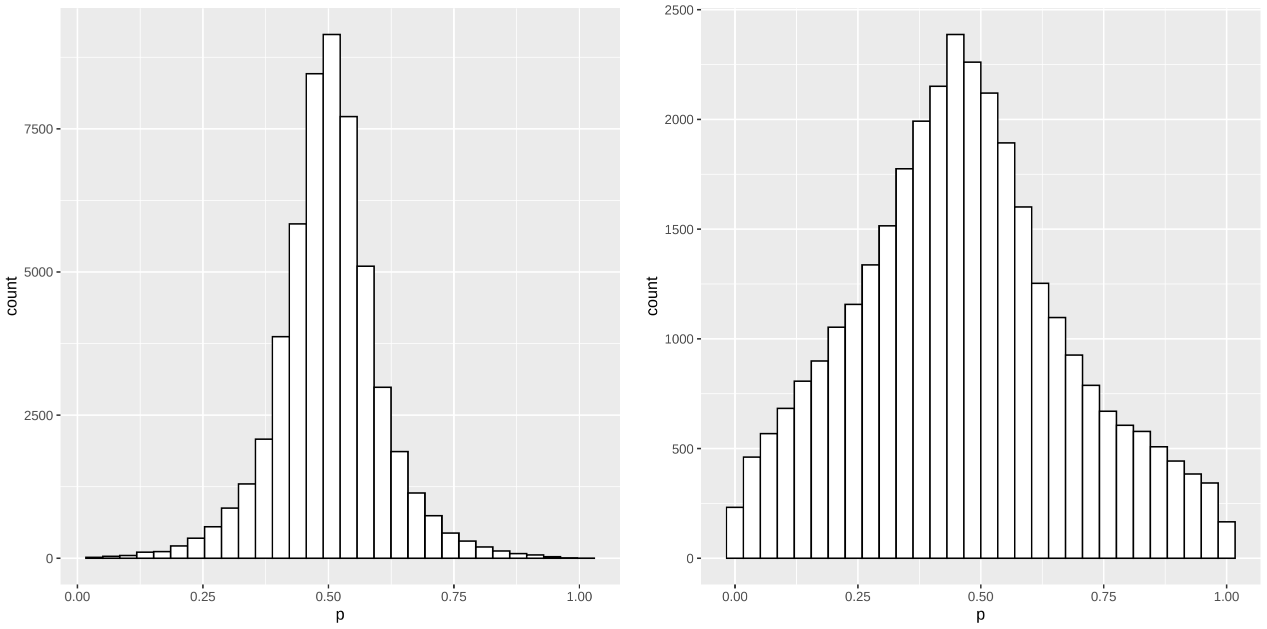


**Supplementary Figure 5:** Histogram of $p$, the probability that group $B$ is up-regulated (while $1 - p$ is the probability that $B$ is down-regulated), in the null simulation studies. Left: bulk simulation; right: single-cell simulation.

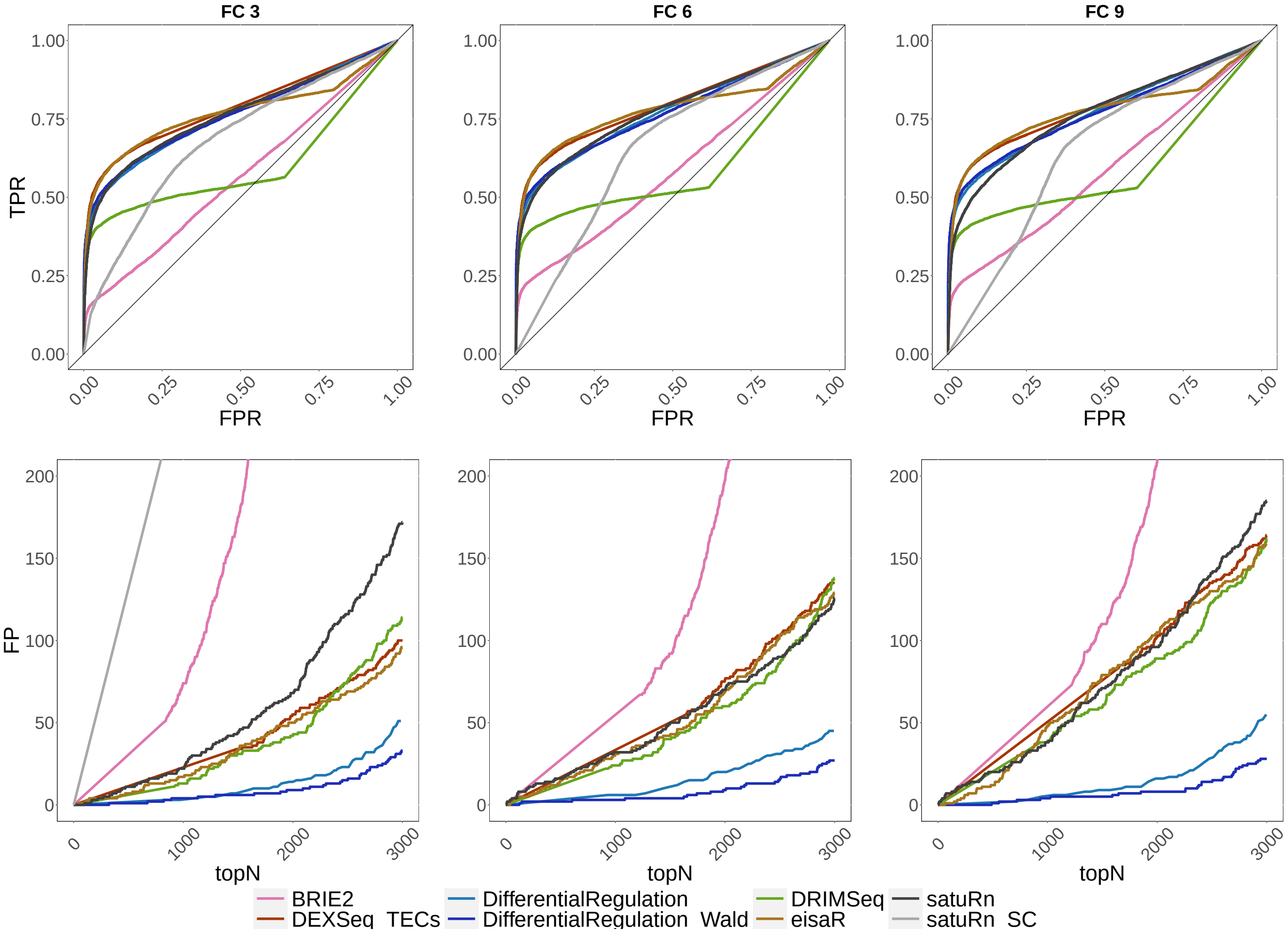


**Supplementary Figure 6:** Single-cell simulation study with DR + DGE. Top row: ROC curves; i.e., false positive rate (FPR) *vs.* true positive rate (TPR). Bottom row: false positive (FP) results among top detections (topN). All simulations have differentially regulated and differentially expressed genes, with average fold-change (FC) of 3 (left panel), 6 (middle panel) or 9 (right panel).

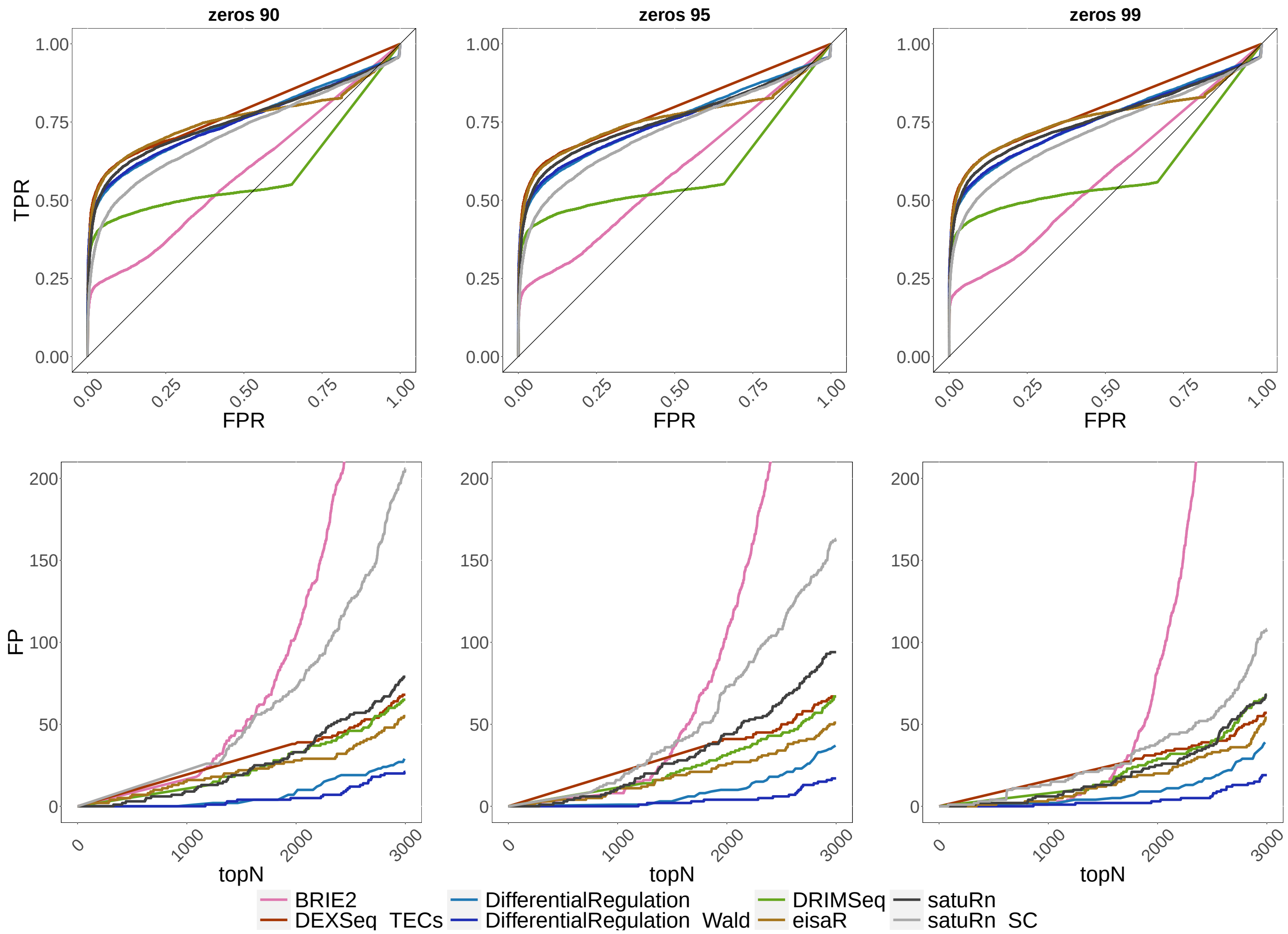


**Supplementary Figure 7:** Single-cell simulation study with various fractions of zero-abundant cells. Top row: ROC curves; i.e., false positive rate (FPR) *vs.* true positive rate (TPR). Bottom row: false positive (FP) results among top detections (topN). All simulations have differentially regulated genes only (no DGE or DAS), with 90 (left column), 95 (middle column) and 99% (right column) of zero cells (i.e., gene expression equal to 0). The average gene abundance is the same across simulations.

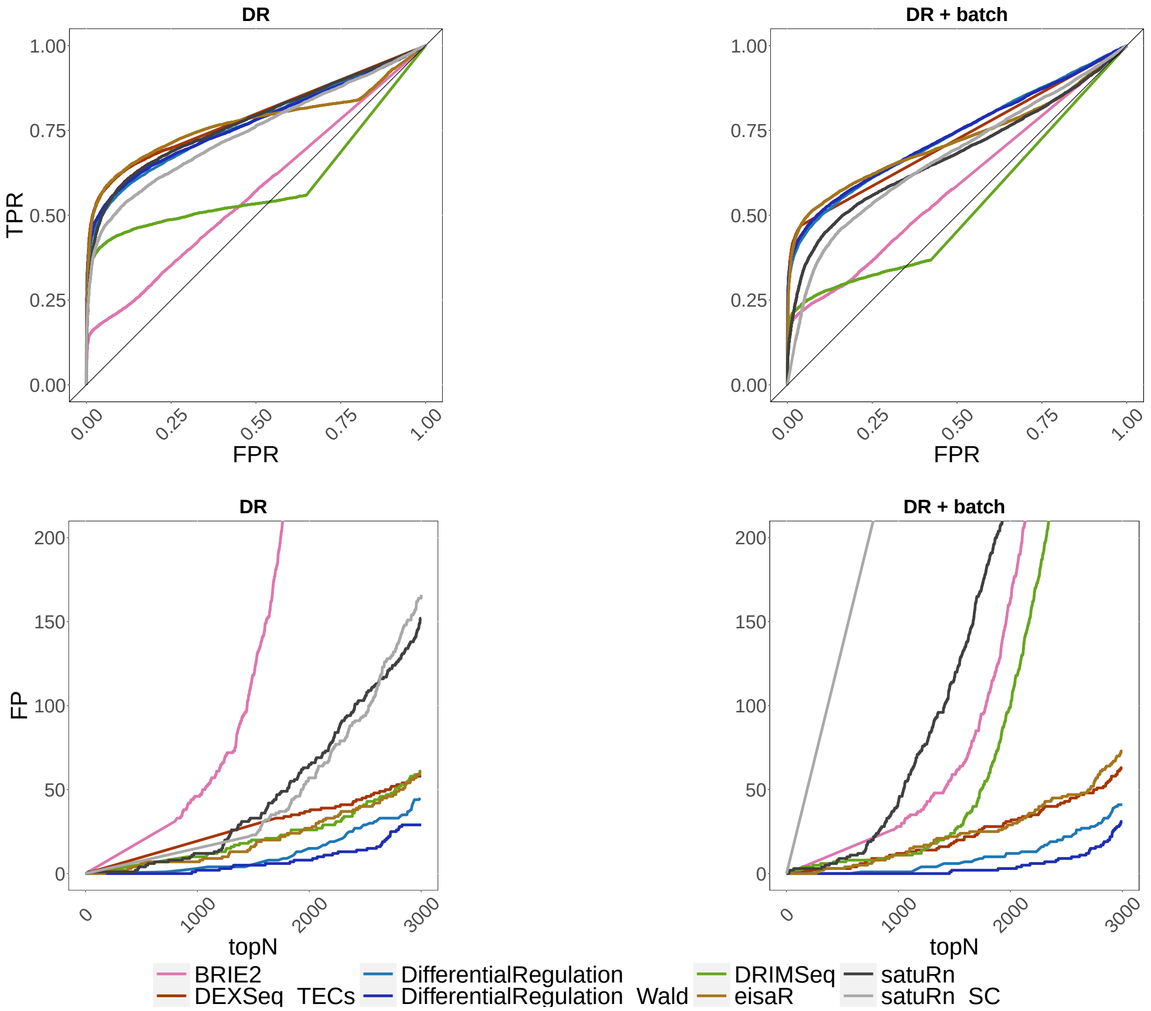


**Supplementary Figure 8:** Results from the single-cell simulations, with and without batch effects. Top row: ROC curves; i.e., false positive rate (FPR) *vs.* true positive rate (TPR). Bottom row: false positive (FP) results among top detections (topN). Left panel (DR): simulation with differential regulation only; right panel (DR + batch): simulation with differential regulation and batch effect.

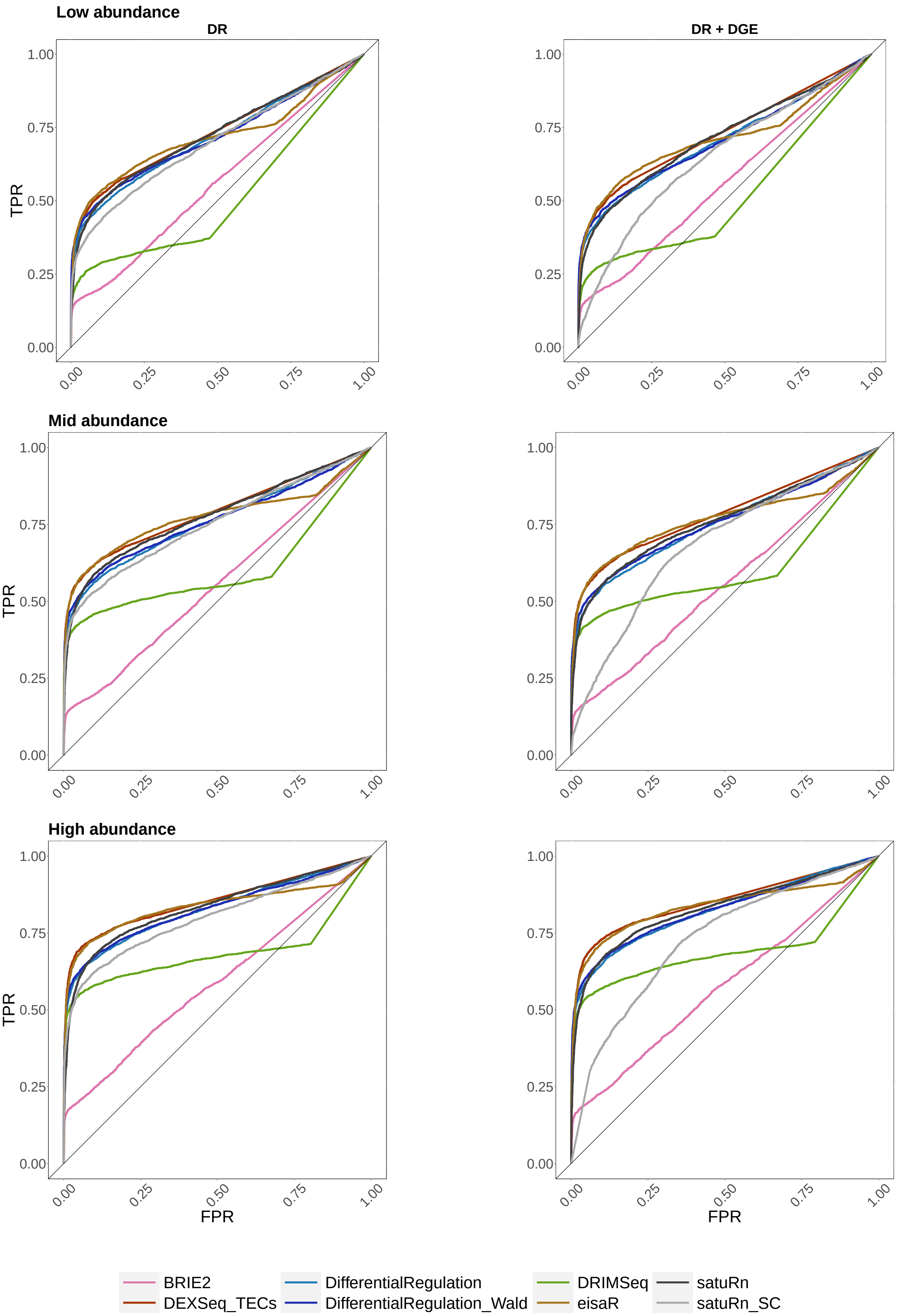


**Supplementary Figure 9:** ROC curves for the main scRNA-seq simulations, stratified by overall transcript abundance: false positive rate (FPR) *vs.* true positive rate (TPR). Top row: low abundance (bottom 33% of abundant transcripts); middle row: medium abundance (middle 33% of abundance); bottom row: high abundance (top 33% of abundant transcripts). Left column: DR only simulation; right column: DR and DGE simulation (average fold-change of 3).

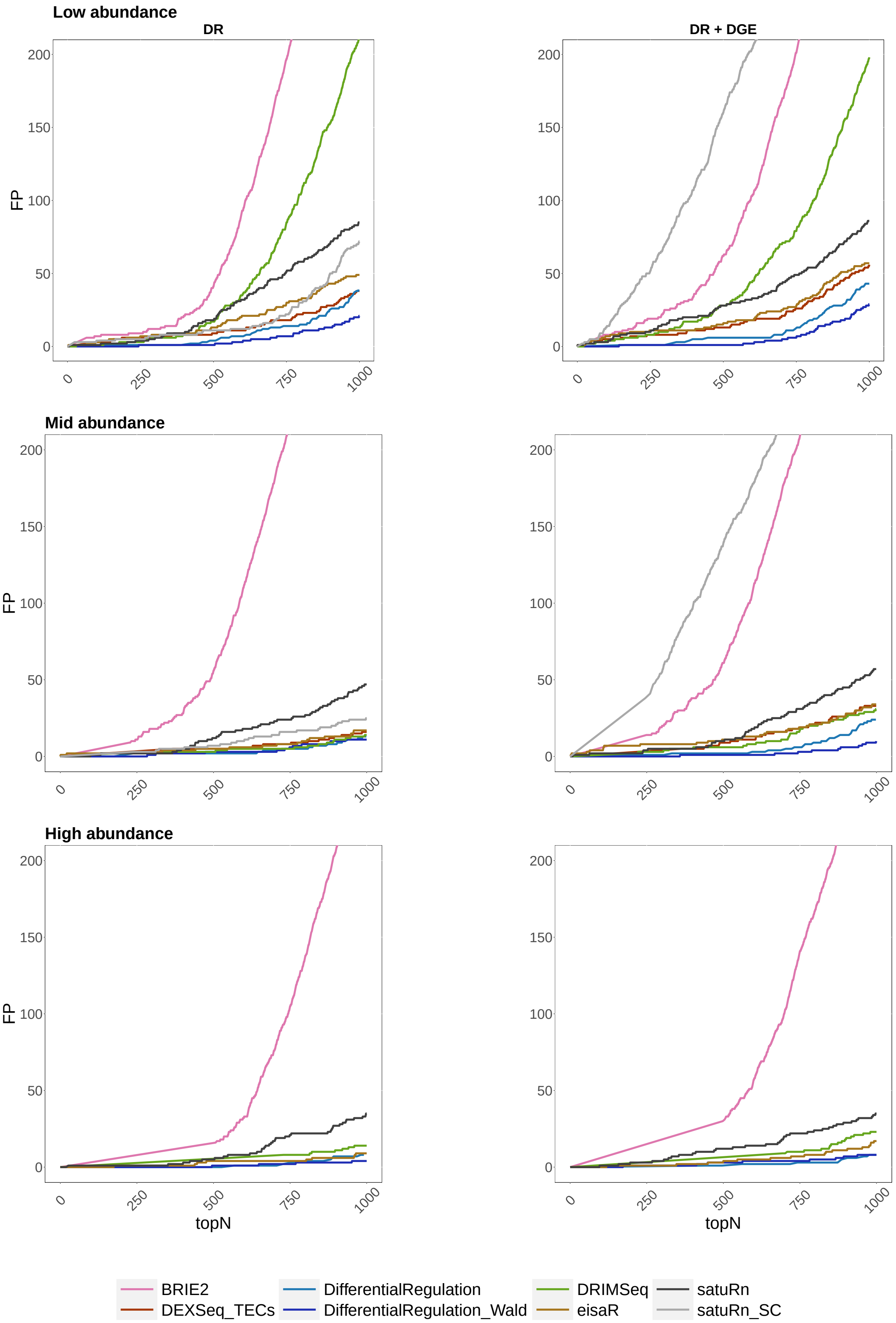


**Supplementary Figure 10:** False positive (FP) results among top detections (topN) for the main scRNA-seq simulations, stratified by overall transcript abundance: false positive rate (FPR) *vs.* true positive rate (TPR). Top row: low abundance (bottom 33% of abundant transcripts); middle row: medium abundance (middle 33% of abundance); bottom row: high abundance (top 33% of abundant transcripts). Left column: DR only simulation; right column: DR and DGE simulation (average fold-change of 3).

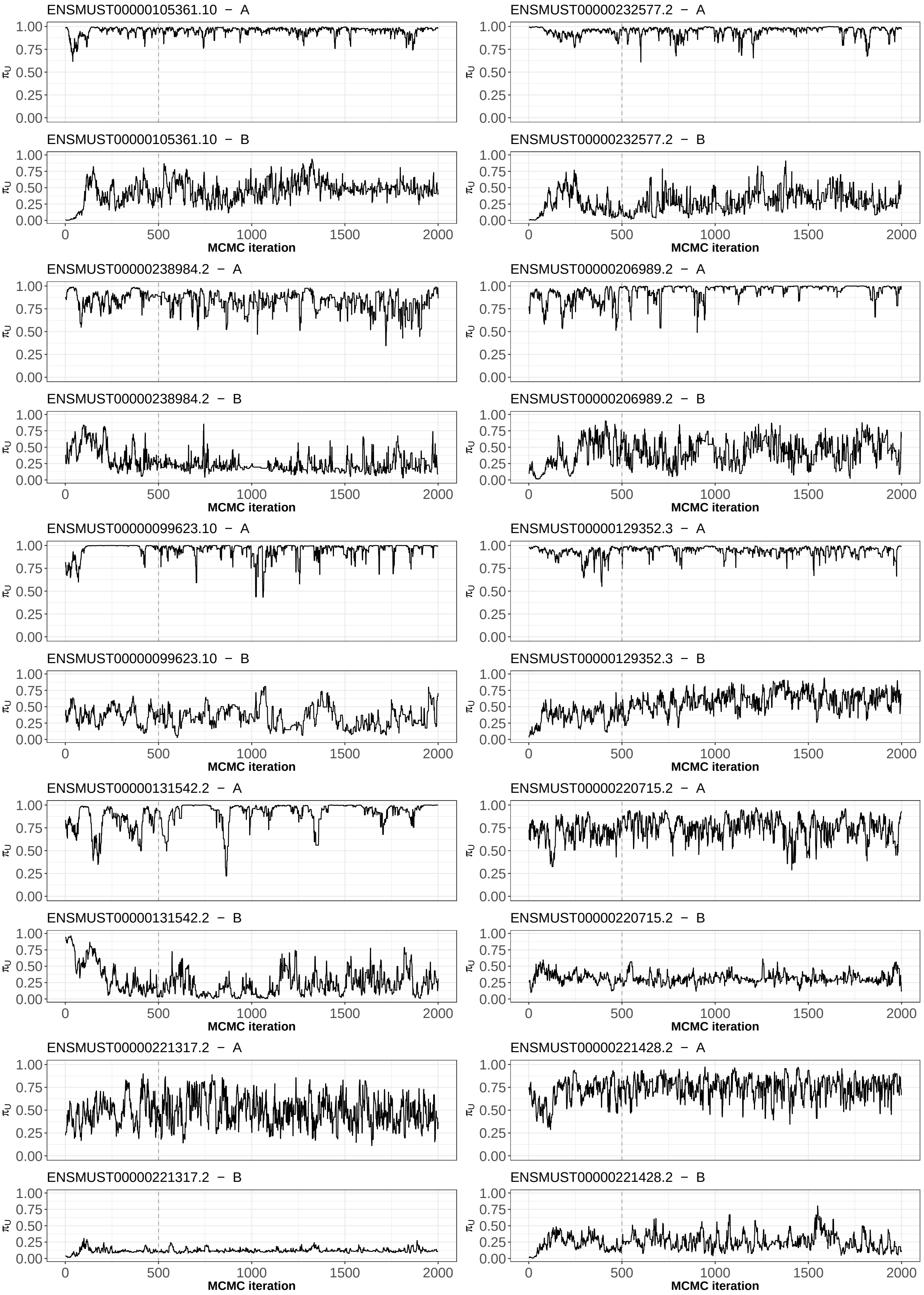


**Supplementary Figure 11:** Traceplot for the posterior distribution of $\tilde{\pi}_U$ (i.e., the key parameter of inference) for the 20 most significant transcripts of the bulk real data analysis (results 1-10). The dashed vertical grey line dentotes the burn-in (excluded from the analyses).

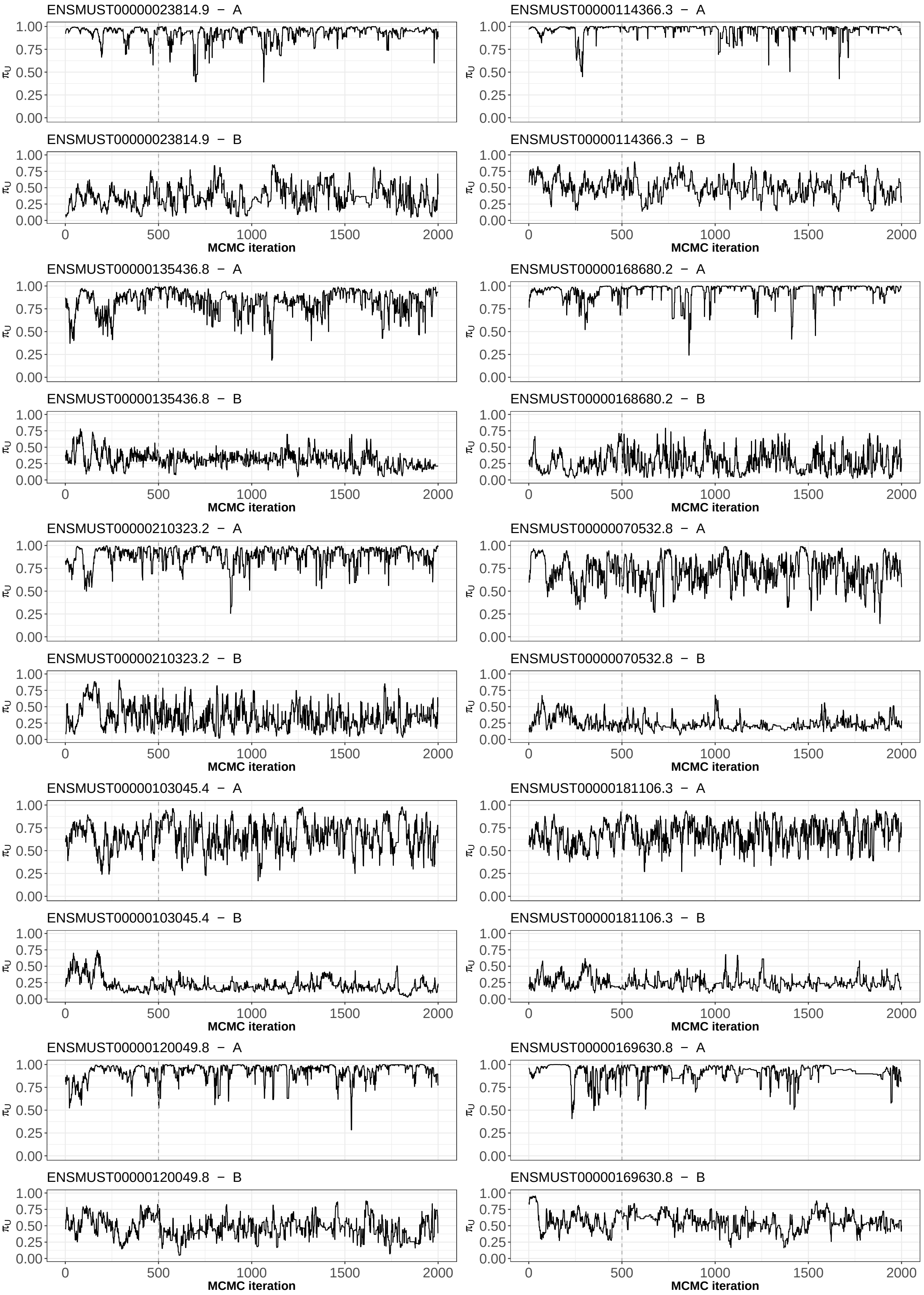


**Supplementary Figure 12:** Traceplot for the posterior distribution of $\tilde{\pi}_U$ (i.e., the key parameter of inference) for the 20 most significant transcripts of the bulk real data analysis (results 11-20). The dashed vertical grey line dentotes the burn-in (excluded from the analyses).

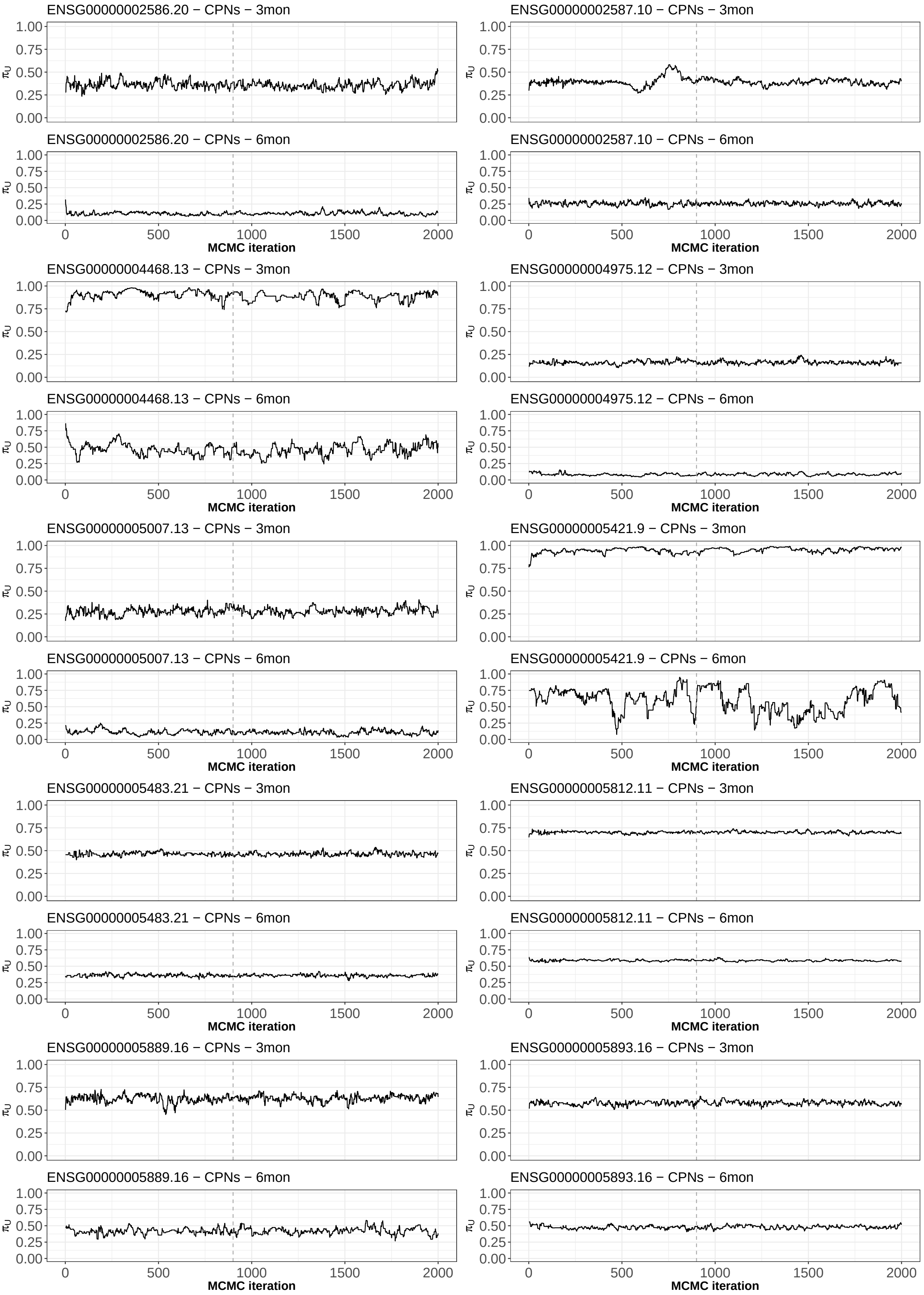


**Supplementary Figure 13:** Traceplot for the posterior distribution of $\tilde{\pi}_U$ (i.e., the key parameter of inference) for the 20 most significant transcripts of the single-cell real data analysis (results 1-10). The dashed vertical grey line dentotes the burn-in (excluded from the analyses).

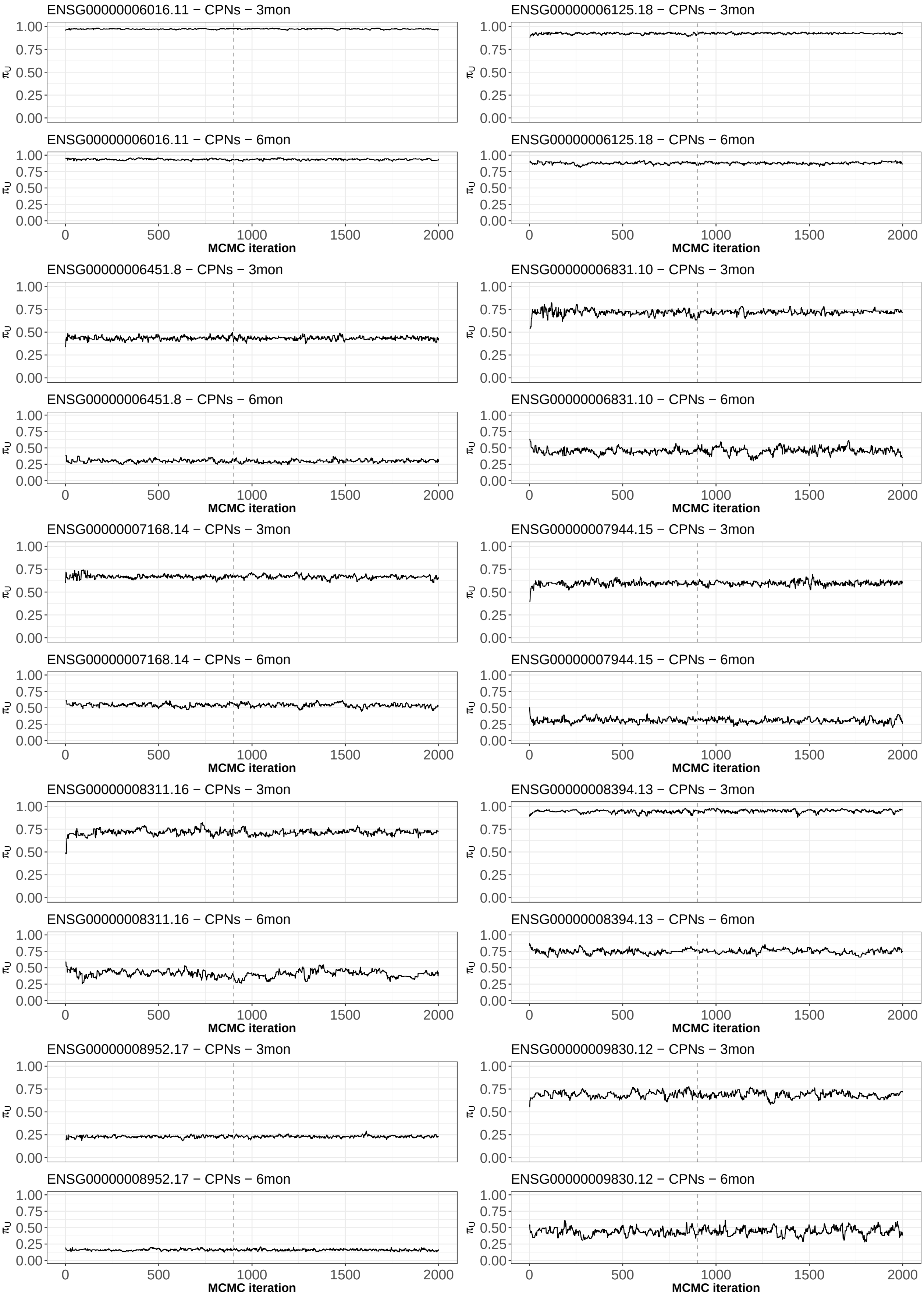


**Supplementary Figure 14:** Traceplot for the posterior distribution of $\tilde{\pi}_U$ (i.e., the key parameter of inference) for the 20 most significant transcripts of the single-cell real data analysis (results 11-20). The dashed vertical grey line dentotes the burn-in (excluded from the analyses).

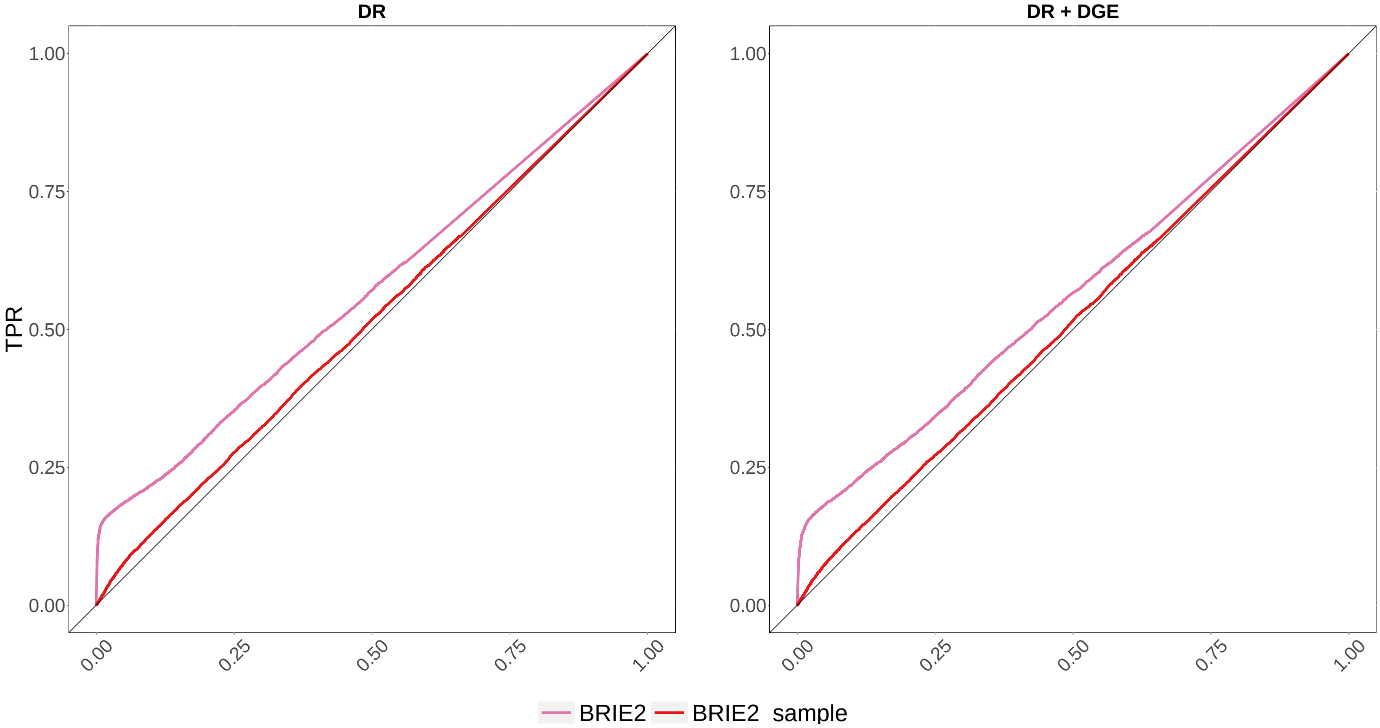


**Supplementary Figure 15:** ROC curves for the main scRNA-seq simulations: false positive rate (FPR) *vs.* true positive rate (TPR). Here, we focus on *BRIE2*, with (*BRIE2_sample*) and without (*BRIE2*) sample as a covaraite in the regression model. Left column: DR only simulation; right column: DR and DGE simulation (average fold-change of 3).
